# Direct phosphorylation and stabilization of HIF-1α by PIM1 kinase drives angiogenesis in solid tumors

**DOI:** 10.1101/2021.03.17.435865

**Authors:** Andrea L. Casillas, Shailender S. Chauhan, Rachel K. Toth, Alva G. Sainz, Amber N. Clements, Corbin C. Jensen, Paul R. Langlais, Cindy K. Miranti, Anne E. Cress, Noel A. Warfel

## Abstract

Angiogenesis is essential for sustained growth of solid tumors. Hypoxia-inducible factor 1 (HIF-1) is a master regulator of angiogenesis and constitutive activation of HIF-1 is frequently observed in human cancers. Thus, understanding mechanisms governing the activation of HIF-1 is critical for successful therapeutic targeting of tumor angiogenesis. Herein, we establish a new regulatory mechanism responsible for the constitutive activation of HIF-1α in cancer, irrespective of oxygen tension. PIM1 kinase directly phosphorylates HIF-1α at threonine 455, a previously uncharacterized site within its oxygen-dependent degradation domain. This phosphorylation event disrupts the ability of prolyl hydroxylases (PHDs) to bind and hydroxylate HIF-1α, interrupting its canonical degradation pathway and promoting constitutive transcription of HIF-1 target genes. Overexpression of PIM1 is sufficient to stabilize HIF-1α in normoxia and stimulate angiogenesis in a HIF-1-dependent manner *in vivo*. CRISPR mutants of HIF-1α (Thr455D) showed increased tumor growth, proliferation and angiogenesis. Moreover, T455D xenograft tumors were refractory to the anti-angiogenic and cytotoxic effects of PIM inhibitors. These data identify a new signaling axis responsible for hypoxia-independent activation of HIF-1 and expand our understanding of the tumorigenic role of PIM1 in solid tumors.

## Introduction

Angiogenesis, or the branching of blood vessels, is a rate-limiting step in the development of solid tumors (Folkman, 1971). Without angiogenesis, solid tumors cannot sustain proliferation due to a lack of oxygen and nutrients (Muthukkaruppan et al., 1982). Hypoxia-inducible factor 1 (HIF-1) is a basic helix-loop-helix-PAS domain transcription factor that is a critical mediator of the cellular response to oxygen deprivation and a key driver of tumor angiogenesis (Semenza, 1999). HIF-1 is a heterodimer that consists of a constitutively expressed subunit, HIF-1β, and HIF-1α, a subunit whose expression is tightly regulated in an oxygen-dependent manner. In the presence of oxygen, cytoplasmic HIF-1α is rapidly hydroxylated on prolines 402 (Pro402) and 564 (Pro564), located within its oxygen-dependent degradation domain (ODDD), by prolyl hydroxylase-domain proteins (PHDs) 1–3 (Bruick and McKnight, 2001; Semenza, 2004). Hydroxylation causes the von Hippel-Lindau (VHL) tumor suppressor protein to recognize HIF-1α and recruit a ubiquitin-protein ligase complex that leads to the ubiquitination and rapid degradation of HIF-1α by the 26S proteasome (Maxwell et al., 1999). In the absence of oxygen, PHDs become inactive and no longer hydroxylate HIF-1α, allowing it to accumulate in the cell and translocate to the nucleus, where it binds to hypoxia-response elements (HREs) to promote the transcription of target genes (Ivan et al., 2001). Therefore, understanding the mechanism by which tumor cells sustain HIF-1α expression is critical for understanding solid tumor growth and identifying new and effective ways to target angiogenesis therapeutically.

Constitutive expression of HIF-1α is common in human cancers, regardless of oxygen tension. Stabilization of HIF-1α in normoxia has been attributed to genetic alterations, such as loss of VHL, as well as transcriptional upregulation due to the activation of oncogenic signaling pathways, such as NF-κB, STAT3, and Sp1. HIF-1α mRNA and protein synthesis are also induced by oncogenic signaling pathways, including PI3K and RAS (Baldewijns et al., 2010; Hua Zhong, 2000; Richard DE, 1999). Post-translational modification also plays a critical role in controlling HIF-1α expression and function. Direct phosphorylation by ERK blocks the nuclear export of HIF-1α and promotes its accumulation in the nucleus, resulting in higher transcriptional activation (Mylonis et al., 2006). It has also been reported that phosphorylation of HIF-1α by various kinases can control its protein stability. Phosphorylation by glycogen synthase kinase 3 and Polo-like kinase 3 promotes HIF-1α degradation (Flügel D, 2007; Isaacs et al., 2002; Xu D, 2010), whereas phosphorylation by CDK1, ATM, and PKA have been reported to stabilize HIF-1α (Bullen et al., 2016; Cam et al., 2010; Warfel et al., 2013). Regardless of the mechanism, stabilization of HIF-1α in normoxia is able to overcome negative regulators, such as Factor inhibiting HIF (FIH-1), and upregulate genes that initiate and sustain angiogenesis during tumor growth (Hartwich et al., 2013). Hence, the identification of novel, oxygen-independent mechanisms that regulate HIF-1α expression is extremely valuable in the effort to understand tumor progression and effectively target angiogenesis as a therapeutic strategy.

The Proviral integration site for Moloney murine leukemia virus (PIM) kinases are a family of serine-threonine kinases that are known to promote tumorigenesis by impacting cell cycle progression, survival, and proliferation (Malone et al., 2020; Nawijn et al., 2011). Pim1 expression is elevated in ~50% of human prostate cancer specimens, particularly in high Gleason grade and aggressive metastatic prostate cancer cases, highlighting its ability to enhance tumorigenesis (Chen et al., 2005; Dhanasekaran et al., 2001). PIM kinases are also elevated in a host of other solid tumors, including colon, breast, and lung cancer, and their overexpression is associated with higher staging, increased metastasis, and diminished overall survival (Braso-Maristany et al., 2017; Chauhan et al., 2020; Gao et al., 2019; Xie Y, 2006; Zhang et al., 2018; Zhao et al., 2017). As a result, several small molecule PIM kinase inhibitors are actively being tested as anti-cancer agents in clinical trials. We recently reported that PIM inhibitors display synergistic anti-tumor effects in combination with anti-angiogenic agents in prostate and colon cancers that was characterized by a dramatic loss of vasculature (Casillas et al., 2018; Malone et al., 2020). Here, we establish PIM1 as an important factor responsible for driving tumor angiogenesis. Mechanistically, we show that PIM1 promotes angiogenesis through a novel signaling axis directly linking PIM1 to HIF-1 via a previously uncharacterized direct phosphorylation event that disrupts the canonical HIF-1α degradation pathway. Our results suggest that the ability of PIM1 to induce angiogenesis and tumor growth is dependent on stabilization of HIF-1 and that the anti-tumor effects of PIM inhibitors are largely due to their anti-angiogenic properties.

## Results

### PIM1 expression is correlated with angiogenesis in human cancers

Tumor vasculature is responsible for providing the nutrients and oxygen required for survival and proliferation, as well as a route for dissemination. We recently discovered that overexpression of PIM1 kinase was sufficient to sustain vasculature during treatment with anti-VEGF therapy (Casillas et al., 2018). Therefore, we sought to determine whether PIM1 expression was correlated with tumor angiogenesis. To this end, we performed immunohistochemistry (IHC) to quantify PIM1 and CD31 (an endothelial cell marker) levels in three prostate cancer tissue microarrays (TMAs) (#2 n = 36, #5 n = 44, and #13 n = 29). These TMAs were obtained from diagnostic cores of radical prostatectomies of patients prior to treatment at the University of Arizona, and samples ranged in severity of disease (Gleason score 6-9). We observed a statistically significant correlation between PIM1 expression and microvessel density (Fig. 1A and B) indicating that PIM1 expression is significantly correlated with vascularization in human tumor samples. To provide further evidence for the association between PIM1 and angiogenesis, we investigated the correlation between *PIM1* and *PECAM1* transcript levels in publicly available data from The Cancer Genome Atlas database. Prostate, colon, and lung cancer all displayed a statistically significant correlation between *PIM1* and *PECAM1* (Fig. S1A). Thus, PIM1 expression is significantly correlated with vascularization in human tumor samples. With the correlative relationship between PIM1 and tumor vasculature established in human tumors, we next sought to determine whether PIM1 is sufficient to promote angiogenesis in vivo using pre-clinical imaging modalities to quantitatively measure changes in vascular perfusion over time. To this end, 1 × 10^6^ PC3 prostate cancer cells were stably transfected with PIM1 or a vector control (VEC) and implanted subcutaneously into the flanks of male NSG mice. Verifying previous results, PIM1-expressing tumors grew significantly faster than wild-type tumors (Fig. 1C, (Casillas et al., 2018)). To assess angiogenesis, a paramagnetic contrast agent (CA) was injected intravenously, and magnetic resonance imaging (MRI) was performed to visualize uptake of the CA in the tumor and reference tissue and generate a time-concentration curve. The resulting curve was fit to a pharmacokinetic model to estimate physiological parameters for the tissue of interest. As the CA passes through the circulation (typically 45–60 s after injection), it is predominantly intravascular, allowing for the evaluation of perfusion (i.e., blood flow per unit volume). During the subsequent 2–15 min, the contrast agent passes into the extravascular space, allowing for measurement of vascular permeability (R_ktrans_) and relative clearance rate. Tumors were imaged once they reached approximately 300 mm^3^, and each cohort was imaged again 7 days later. To compare angiogenesis, DCE-MRI data from size-matched PC3/VEC and PC3/PIM1 tumors were analyzed using the Linear Reference Region Model, which has superior resolution to current methods (Cardenas-Rodriguez et al., 2013). The resulting time-concentration curve revealed a dramatic increase in the uptake of the CA in PIM1-expressing tumors compared to control tumors, indicating that PIM1 expression significantly increases perfusion (Fig. 1D). Furthermore, the average vascular permeability (R_ktrans_) was significantly greater in PIM1-overexpressing tumors than in control tumors (Fig. 1E). Taken together, these analyses indicate that PIM1-overexpressing tumors have more blood flow and a greater number of mature vessels compared to VEC. This correlation was confirmed with endpoint staining of CD31, which demonstrated that microvessel density was significantly higher in PIM1 tumors than in control tumors (Fig. S1B and C). Taken together, these data demonstrate that PIM1 expression is sufficient to enhance tumor angiogenesis.

**Figure 1.**
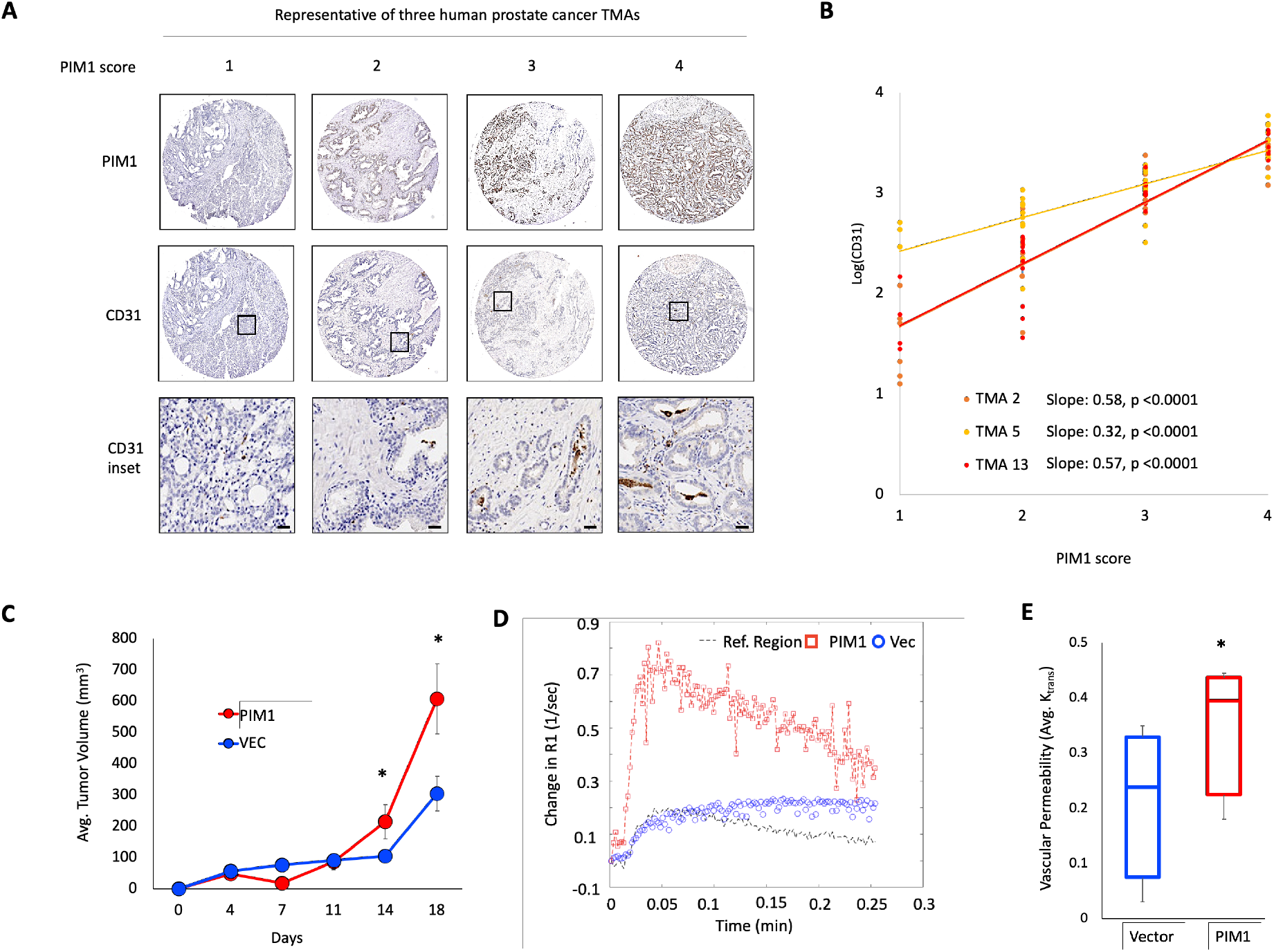
PIM1 correlates with angiogenesis in human cancer samples. A) Representative PIM1 and CD31 staining of three human prostate cancer TMAs, n=109. B) Scoring of PIM1 and CD31 were graphed to determine the association between PIM1 and vasculature by Pearson correlation in each of the indicated TMAs. C) Mice (n=4/group) were injected with PC3-Vec or PC3-PIM1 cells and tumor volume was measured via caliper over time. D) Representative DCE-MRI trace from size-matched PC3 PIM1 and PC3-Vec tumors and E) average vascular perfusion (K_trans_). Scale bar = 50 μm. *p < 0.05, error bars = SEM.

### PIM1 induces angiogenesis in a HIF-1-dependent manner

HIF-1 is a transcription factor that is a master regulator of angiogenesis, and we previously reported that PIM inhibitors can reduce the levels of HIF-1α (Casillas et al., 2018). Therefore, we sought to determine whether the pro-angiogenic effect of PIM1 is dependent on HIF-1. To determine whether HIF-1 activation is required for the pro-angiogenic effect of PIM1, HIF-1/2α were knocked down using siRNA in PC3 cells stably expressing a doxycycline (Dox)-inducible lentiviral vector encoding PIM1 (Dox-PIM1) [25], and in vitro angiogenesis assays were performed. Forty-eight hours after knockdown, Dox-PIM1 cells were treated with Dox, and cell lysates and conditioned media (CM) were collected after 24 h. Then, human umbilical vein endothelial cells (HUVECs) were suspended in CM from each experimental condition, plated on reduced growth factor basement membrane, and tube formation was assessed over time. Immunoblotting verified that PIM1 stabilized HIF-1/2α, and siRNA effectively reduced HIF-1/2 α levels (Fig. 2A). Fluorescence images of calcein AM-stained endothelial cells were acquired over 6 h, which corresponded with maximal tube formation (Fig. 2B). CM from Dox-PIM1 cells substantially enhanced tube formation compared to CM from Dox-VEC cells. Strikingly, the pro-angiogenic effect of PIM1 expression was abolished in cells lacking HIF-1/2α. Image analysis verified that PIM1 overexpression significantly increased the average tube length and number of branch points, and HIF-1/2α knockdown restored tube formation to basal levels (Fig. 2C and D). To confirm that PIM1 expression impacts the release of proangiogenic factors from tumor cells through the upregulation of HIF-1 target genes, we measured the amount of vascular endothelial growth factor (VEGF) in the CM from each sample. Conditioned media from PIM1-expressing cells contained nearly 5-fold more VEGF than control cells, and knockdown of HIF-1/2α restored VEGF levels to basal levels (Fig. 2E). We further generated RKO colon cancer cell lines stably overexpressing PIM1 or a vector control in combination with stable knockdown of HIF-1α (Fig. S2A and B). In vitro angiogenesis assays using CM from each cell line demonstrated that PIM1 expression significantly increased mean tube length and total branch points compared to vector control, whereas PIM1 was unable to induce tube formation in RKO shHIF-1α cells (Fig. S2C-E). To show that the pro-angiogenic effect of PIM1 requires HIF-1α in vivo, 5 × 10^6^ RKO cells were injected subcutaneously into the flanks of twelve (n=3/group) SCID mice, and tumor growth was measured over time and angiogenesis was assessed at several time points by *in vivo* imaging. As expected, PIM1-overexpressing tumors grew significantly faster than controls (Fig. 2F). Interestingly, knockdown of HIF-1α abolished the growth advantage of PIM1 overexpression (Fig. 2F). To monitor angiogenesis, mice from each group were injected with Angiosense 750EX, a fluorescent probe that remains localized in the vasculature to allow for *in vivo* imaging of angiogenesis. To calculate the relative amount of functional vasculature in each cohort, Angiosense signal was normalized to differences in tumor volume to obtain a vascular index for each group. RKO tumors expressing PIM1 were significantly more vascular than the parental line, whereas no increase in angiogenesis was observed with PIM1 overexpression in tumors lacking HIF-1α (Fig. 2G). CD31 staining of tumors from each group confirmed that the pro-angiogenic effect of PIM1 was lost in tumors with knockdown of HIF-1α (Fig. 2H and I). Similarly, the expression of HIF-1 target genes (VEGF and HK2) was increased by PIM1 expression *in vivo*, but not in HIF-1α knockdown tumor tissue (Fig. S2F). Moreover, the growth advantage associated with PIM1 overexpression can be attributed to increased survival, not proliferation, as PIM1 significantly reduced cell death (CC3 staining) but did not change Ki67 staining (Fig. 2H). Notably, tumors lacking HIF-1α showed significantly higher death compared to control, regardless of PIM1 overexpression (Fig. 2I). Taken together, these data indicate that expression of PIM1 promotes angiogenesis in a HIF-1-dependent manner and suggests that the pro-tumorigenic effect of PIM1 can largely be attributed to its ability to promote angiogenesis and prevent cell death.

**Figure 2.**
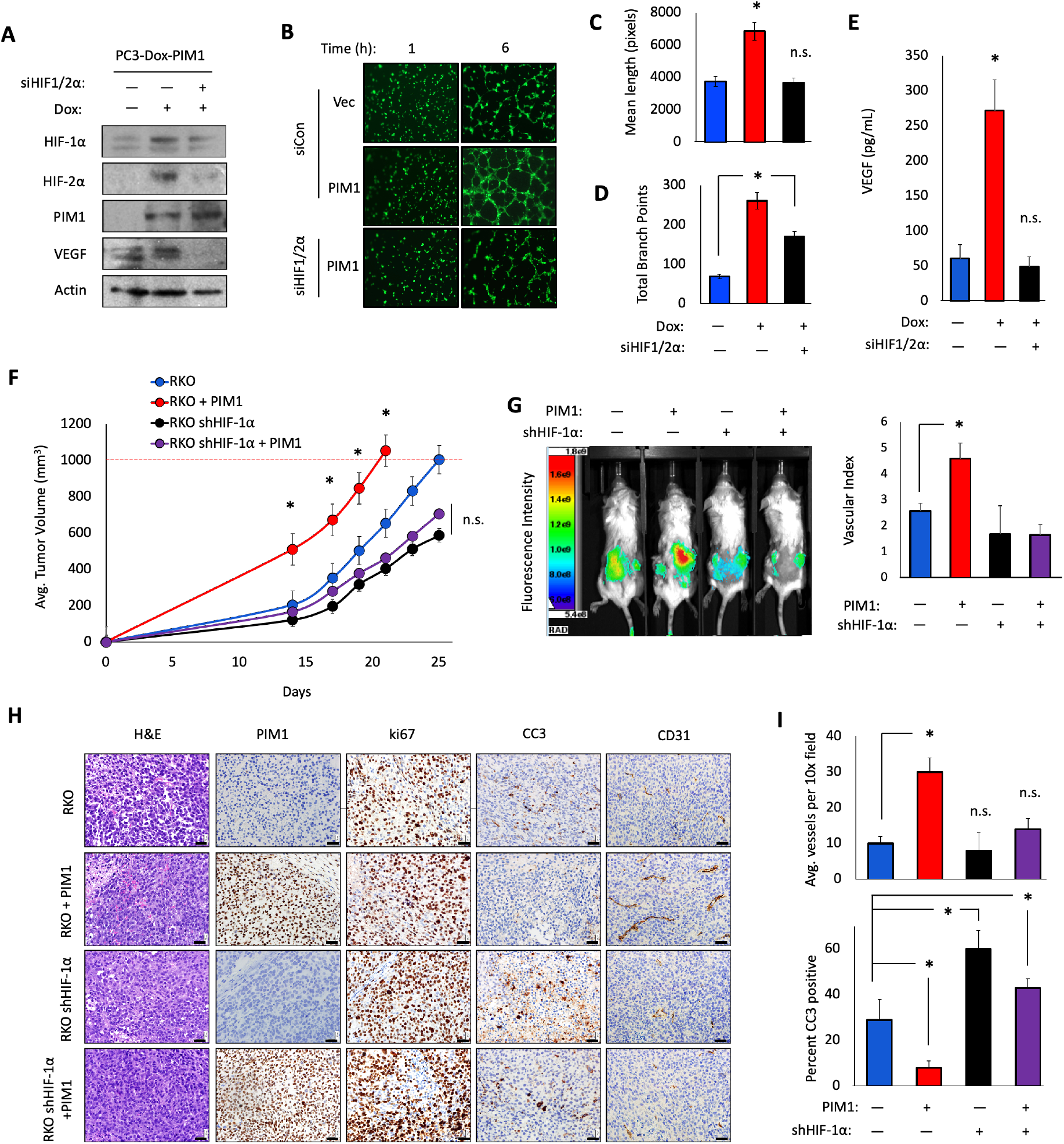
PIM1 induces angiogenesis in vivo and in vitro. A) Dox-PIM1 PC3 cells were transfected with siHIF-1/2α prior to treatment with dox for 24 h, and lysates were collected for immunoblotting and conditioned media (CM) was harvested for *in vitro* angiogenesis assay. B) Representative images of tube formation at 1 and 6 hours after plating HUVEC cells in CM. C) Quantification of mean tube length and D) total branch points as measured with Image J Angiogenesis Analyzer plug-in. E) VEGF-A levels in CM from the indicated conditions were measured by ELISA. F) Twelve mice (n=3/group) were injected with the indicated RKO cell lines, and tumor volume was measured over time. G) Mice were injected with 2 nmol of Angiosense 750EX 24 hours prior to imaging for fluorescence intensity. Vascular index was calculated by normalizing the bioluminescence signal to tumor volume. H) Tumors derived from each cell line were harvested and immunostained with CD31, PIM1, HIF-1α, and CC3. I) Quantification of IHC. Scale bar = 50 μm. *p < 0.05, n.s. = not significant, error bars = SEM.

### PIM1 promotes pro-angiogenic gene expression through HIF-1

Because the pro-angiogenic effect of PIM1 is dependent on HIF-1α, we hypothesized that elevated PIM1 expression is sufficient to activate HIF-1 in the absence of hypoxia. Immunoblotting was used to evaluate HIF-1α protein levels in control and PIM1-overexpressing colon (RKO), prostate (PC3), and lung (A549) cell lines. Strikingly, PIM1 expression was sufficient to stabilize HIF-1α in all cell lines tested (Fig. 3A-C). Importantly, treatment with chemically distinct pan-PIM kinase inhibitors (PIM447 and AZD1208) blocked the ability of PIM1 to increase HIF-1α protein levels, demonstrating that these effects are not due to the chemical itself (Fig. 3A and C). To ensure that the levels of HIF-1α observed after PIM1 induction were sufficient to activate HIF-dependent transcription, Dox-VEC or Dox-PIM1 cells were co-transfected with Renilla-luciferase and a previously described HIF-1 reporter that drives luciferase expression (HRE-Luc) (Rapisarda et al., 2002), treated for 24 h with Dox to induce PIM1, and then treated with DMSO or AZD1208 for 4 h. To account for increased cell growth and death due to PIM1 overexpression and PIM inhibitor treatment, respectively, the HRE-Luc signal was normalized to Renilla-Luc levels. PIM1 expression increased HIF-1 activity by approximately 2-fold in normoxia, and this effect was reversed by treatment with the PIM inhibitor (Fig. 3D). To assess the effect of PIM1 on HIF-1 target genes, we used a semi-high throughput qPCR assay to measure a panel of 84 hypoxia-inducible genes (Qiagen RT Profiler). PC3 Dox-PIM1 cells were cultured in normoxic conditions with or without 20 ng/mL Dox for 24 h to induce PIM1 and treated ± AZD1208 for 8 h, at which point mRNA was collected for subsequent gene expression analysis. PIM1 expression altered the transcript levels of several classes of hypoxia-responsive genes, including critical mediators of angiogenesis, proliferation, and apoptosis (Fig. 3E). To identify genes whose induction was specific to PIM1, we focused on genes that were upregulated at least 3-fold by PIM1 expression in normoxia and significantly reduced by treatment with AZD1208. Of the eleven genes that fit these criteria, seven are known to promote angiogenesis, and all are established targets of HIF-1 (Fig. 3F). We validated that PIM1 increased the expression of several well-known HIF-1 target genes (*VEGF-A*, *ANGPT4*, and *HK2*) by qRT-PCR, and treatment with PIM447 restored the expression of each to basal levels (Fig. 3G). To confirm that PIM1 alters gene expression in a HIF-1-dependent manner, we assessed the expression of the same set of HIF-1 target genes in the previously described RKO cell line with stable knockdown of HIF-1α (Fig. S2C and D). PIM1 overexpression significantly increased the transcript level of all three genes compared to control cells, whereas no increase was observed in RKO cells lacking HIF-1α (Fig. 3H). Thus, PIM1 expression is sufficient to stabilize HIF-1/2α in normoxic conditions and increase the transcription of HIF-1 target genes.

**Figure 3.**
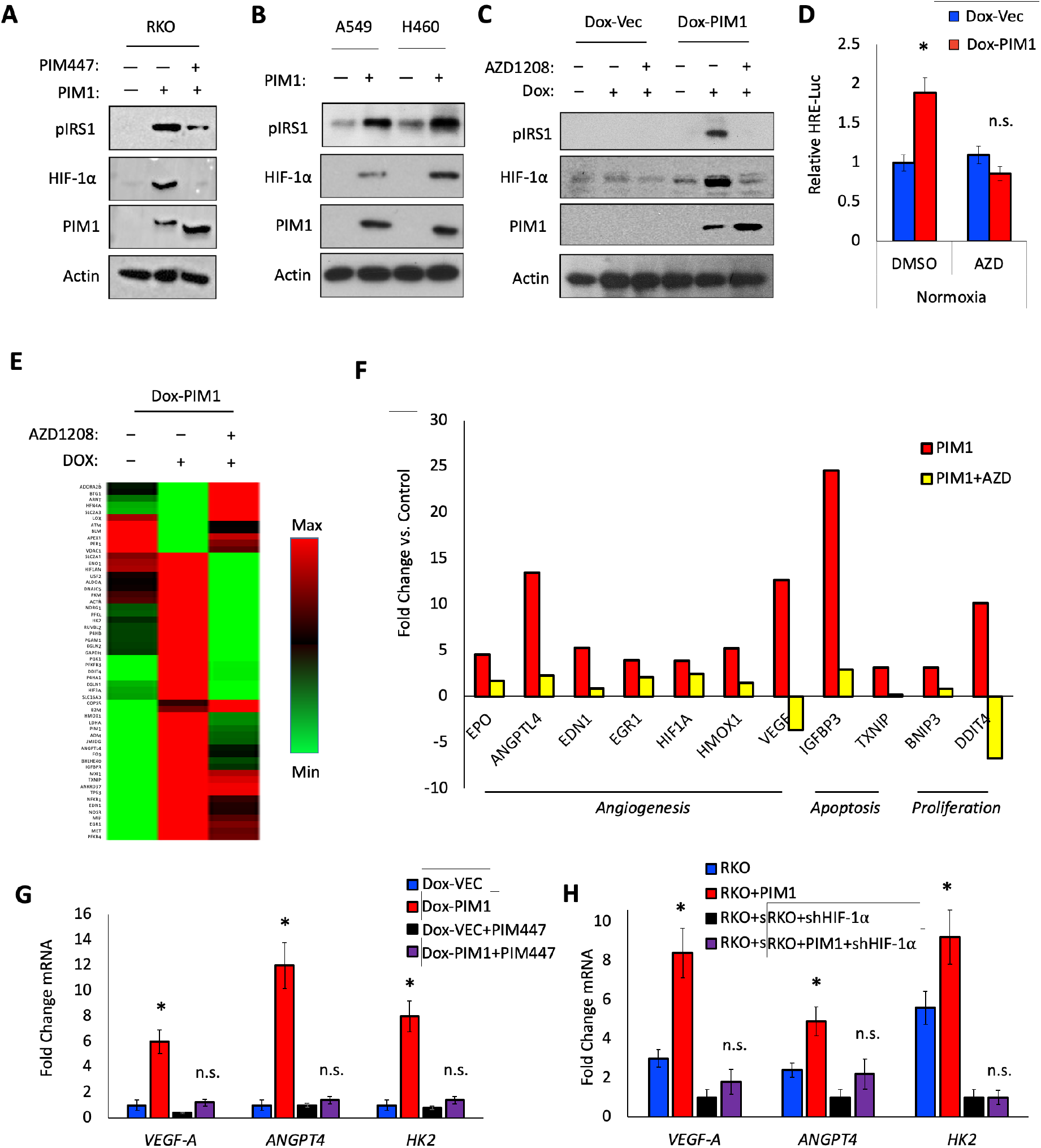
PIM1 is sufficient to stabilize HIF-1α and activate HIF-1 in normoxia. A) RKO colon cancer cells ± PIM1 were treated with DMSO or PIM447 (1 μM) for 6 h. B) A549 and H460 lung cancer cells were stably infected with lentiviral constructs expressing Vector or PIM1. C) Dox-Vec or Dox-PIM1 PC3 cells were treated with dox for 24 h prior to DMSO or AZD1208 (3 μM) for 6 h. D) Dox-PIM1 expressing HRE-Luc were treated with Dox for 24 h prior to DMSO or AZD1208 for 6 h, and bioluminescence was measured. E and F) Dox-PIM1 cells were treated with Dox for 24 h prior to DMSO or AZD1208 for 6 h and RNA was harvested to measure the expression of hypoxia-inducible genes. F) HIF-1 target genes upregulated by 3-fold by PIM1 and reduced by AZD1208. G) Dox-PIM1 cells were treated with Dox for 24 h prior to DMSO or AZD1208 for 6 h and RNA was harvested to measure gene expression by qRT-PCR. H) RNA was harvested from the indicated cell lines and gene expression was measured by qRT-PCR. *p < 0.05, n.s. = not significant, error bars = SEM.

### PIM1 phosphorylates HIF-1α at Threonine 455

Next, we sought to determine the mechanism by which PIM1 stabilizes HIF-1α. Because PIM1 is a serine-threonine kinase, we hypothesized that PIM1 may directly phosphorylate HIF-1α to alter its protein stability. First questioned whether these proteins interact in cells. To this end, HA-PIM1 was overexpressed in 293T cells and treated for 6 h with MG132 in normoxia or hypoxia to stabilize HIF-1α prior to harvest. PIM1 was immunoprecipitated using the HA-tag and HIF-1α binding was assessed by western blotting. PIM1 pulled down both endogenous HIF-1α in both normoxia and hypoxia, demonstrating that these proteins complex in cells (Fig 4A). Next, *in vitro* kinase assays were performed using recombinant PIM1 and HIF-1α. Autoradiography revealed that PIM1 is able to phosphorylate HIF-1α, and phosphorylation was lost in the presence of a PIM inhibitor (Fig. 4B). To identify PIM-mediated phosphorylation sites, HIF-1α was isolated and mass spectrometry analysis was performed to identify post-translational modifications. PIM1 phosphorylated HIF-1α at two sites *in vitro*. The first, Ser643, has been previously described as an ERK target site that enhances the nuclear localization of HIF-1α but does not alter its protein stability (Mylonis et al., 2006). The second site, Thr455, is a previously uncharacterized site located within the ODDD of HIF-1α between Pro402 and Pro564, which are hydroxylated by PHDs as a signal initiating the proteasomal degradation of HIF-1α (Fig. 4B). Notably, Thr455 is evolutionarily conserved among mammals, suggesting its importance as a regulatory site (Fig. 4C). Based on its localization within the ODDD, we chose to further investigate the effect of Thr455 phosphorylation on HIF-1α stability. To verify that this site is phosphorylated in cells, PIM1 and HA-HIF-1α were co-transfected into 293T cells, HIF-1α was immunoprecipitated, and mass spectrometry was used to detect post-translational modifications. HIF-1α was robustly phosphorylated at Thr455 in cells expressing PIM1 (Fig. S3A). Next, we generated a phospho-specific antibody against HIF-1α Thr455. To verify the specificity of this antibody, we used site-directed mutagenesis to create a T455A mutant of HIF-1α. Wild-type or T455A HA-HIF-1α were immunoprecipitated from cells and incubated with recombinant PIM1 in the same conditions used for *in vitro* kinase assays. Wild-type HIF-1α displayed robust phosphorylation at Thr455 by PIM1, which was blocked by AZD1208, whereas the T455A mutant was not recognized by the phospho-antibody (Fig. 4E). Additionally, staining of tumor sections from Fig 2J-K showed increased phosphorylation of HIF-1α (T455) in tumor tissue over expressing PIM1 compared to control, and no staining was observed in HIF-1α knockdown tumors (Fig. S3B and C). These results confirm that PIM1 directly phosphorylates HIF-1α at T455 and our antibody recognizes phosphorylation specific to this site. To further confirm that PIM1 phosphorylates HIF-1α in cells, HEK293T cells were transfected with vector, HA-PIM1, or a kinase-dead PIM1 (K67M), and total and phospho-HIF-1α (T455) were assessed by immunoblotting. At the protein level, HIF-1α was only upregulated and phosphorylated in cells expressing kinase-active PIM1 (Fig. 4F, lanes 1-3). To further confirm that Thr455 phosphorylation in PIM1-expressing cells was not solely due to the increased abundance of HIF-1α, 293T cells were treated with a proteasome inhibitor (MG-132) for 4 h to stabilize HIF-1α in normoxia. In this context, phosphorylation of Thr455 was observed in basal conditions and significantly increased upon overexpression of PIM1. Alternatively, expression of kinase-dead PIM1 actually reduced Thr455 phosphorylation below basal levels in MG132 treated cells, suggesting that this construct acts as a dominant negative (Fig. 4F, lanes 4-6). To ensure that Thr455 phosphorylation correlated with changes in endogenous PIM1 expression, we cultured C42-B, DU145, and PC3 prostate cancer cell lines in normoxia or hypoxia (1.0% O_2_) for 4 h, which is known to increase PIM1 levels (Warfel et al., 2016b). As expected, total HIF-1α levels were significantly increased by hypoxia. Interestingly, PIM1 levels were markedly increased by hypoxia in DU145 and PC3 cells, but not did not increase in C42-B cells. Strikingly, Thr455 levels were strongly induced only in the DU145 and PC3 cells in hypoxia, providing further evidence that PIM1 controls this site (Fig. 4G). Next, we altered PIM expression and activity in several cancer cell lines to ensure that Thr455 phosphorylation was universally observed in cancer cell lines in which we established that PIM1 increases HIF-1α. PIM1 overexpression increased Thr455 phosphorylation in RKO colon cancer cells, and this effect was reduced upon knockdown of HIF-1α, demonstrating the specificity of this antibody (Fig. S3D). Similarly, induction of PIM1 in PC3 Dox-PIM1 cells increased Thr455 phosphorylation, and co-treatment with PIM447 blocked the induction of phospho-T455 and total HIF-1α (Fig. S3E). In addition, A549 and H460 lung cancer cell lines overexpressing vector or PIM1 were treated with MG132 for 4 h to stabilize HIF-1α and normalize levels prior to assessing Thr455 phosphorylation. PIM1 overexpression dramatically induced Thr455 phosphorylation in lung cancer cell lines (Fig. S3F). Taken together, these studies establish that PIM1 directly phosphorylates HIF-1α at Thr455.

**Figure 4.**
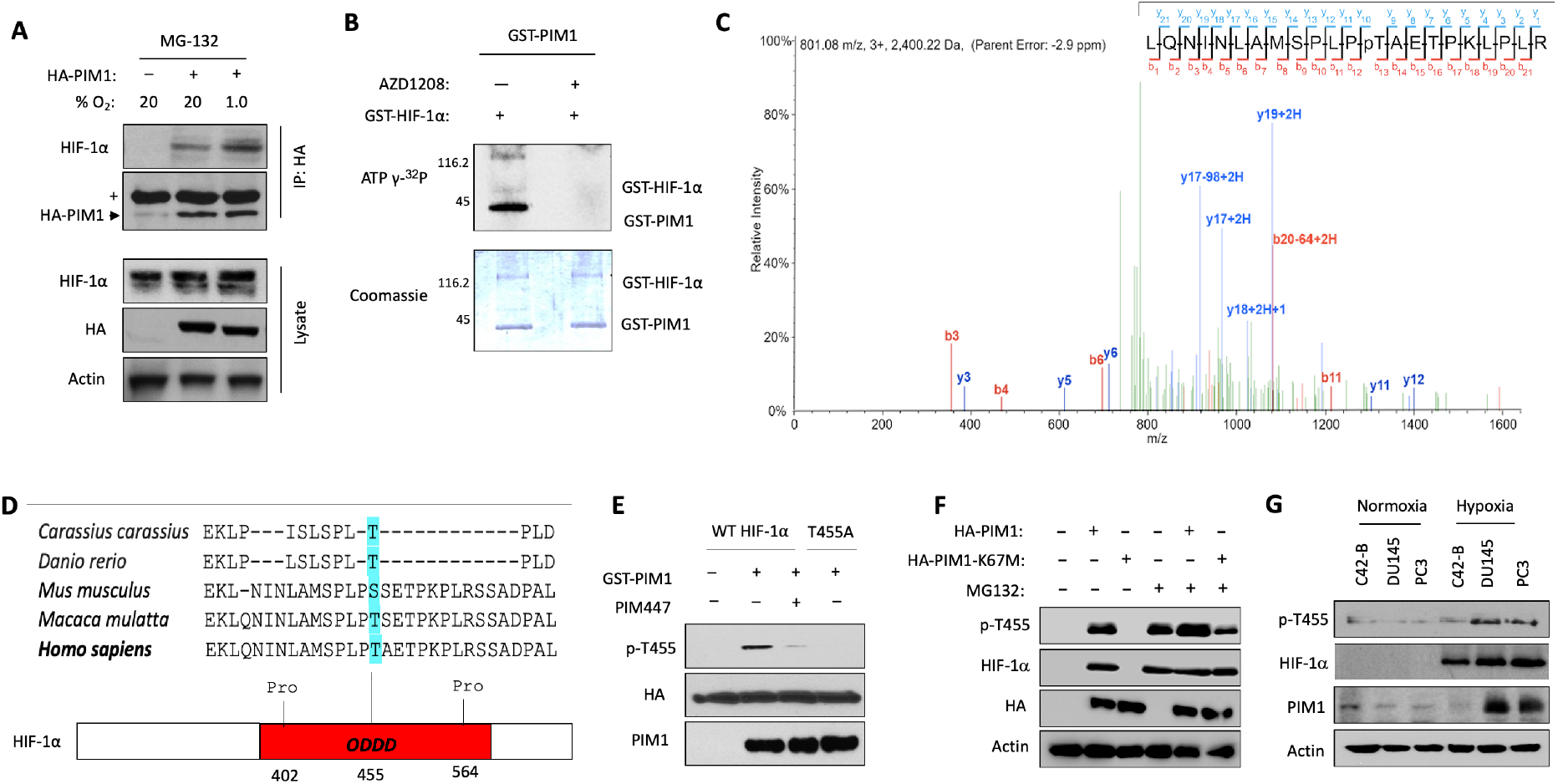
PIM1 phosphorylates HIF-1α at Thr455. A) 293T cells were transfected with HA-PIM1 and with MG-132 in normoxia or hypoxia for 6 h. HA-PIM1 was immunoprecipitated and interaction with endogenous HIF-1α was assessed by western blotting. (+ = Heavy chain IgG). B) Images of Coomassie and autoradiography of *in vitro* kinase assays using recombinant PIM1 and HIF-1α. C) Spectra from mass spectrometry analysis of HIF-1α from *in vitro* kinase assay showing phosphorylation of HIF-1α at Thr455 by PIM1. D) Sequence alignment across species and schematic of HIF-1α ODDD (Thr455 highlighted in blue) E) Immunoblot of *in vitro* kinase assay combining recombinant PIM1 with immunoprecipitated HIF-1α constructs. F) 293T cells were transfected with WT HA-PIM1 or kinase-dead HA-PIM1-K67M and treated with DMSO or MG132 (10 μM) for 3 h. G) The indicated prostate cancer cell lines were cultured in normoxia or hypoxia for 4 h.

### PIM-mediated phosphorylation of HIF-1α at Thr455 increases its protein stability

Next, we characterized the effect of Thr455 phosphorylation on HIF-1α protein levels. Wild-type and RKO cells stably expressing PIM1 were cultured in hypoxia (1% O_2_) for 4 hours to stabilize HIF-1α and then returned to normoxia (20% O_2_), and lysates were collected over a 30-min time course. The half-life of HIF-1α was significantly longer in PIM1-expressing cells than in wild-type cells (30.1 ± 1.2 vs. 9.8 ± 0.5 mins) (Fig. 5A). To directly assess the effect of Thr455 phosphorylation on HIF-1α protein stability, we used site-directed mutagenesis to generate HIF-1α T455D (phosphomimetic) and T455A (phospho-null) constructs. Following transfection of WT, T455D, or T455A HIF-1α, HEK293T cells were treated with cycloheximide and lysates were collected over time to determine the rate of protein degradation. The half-life of the phospho-null mutant (T455A) was significantly shorter than that of WT HIF-1α (1.4 ± 0.2 vs. 2.1 ± 0.2 h), whereas the half-life of the phospho-mimetic (T455D) was significantly longer, showing little degradation over the 4-h time course (Fig. 5B). Because HIF-1α is primarily degraded by the proteasome in normoxic conditions, we tested whether PIM1 decreased HIF-1α ubiquitination. HEK293T cells stably expressing PIM1 or a vector control were transfected with HA-HIF-1α and treated with DMSO or PIM447 for 30 min, followed by MG-132 treatment for 4 h to allow for accumulation of ubiquitinated HIF-1α. HIF-1α was immunoprecipitated and ubiquitin was detected by immunoblotting. PIM1 expression decreased the amount of ubiquitin bound to HIF-1α by approximately 3-fold, whereas PIM inhibition significantly increased the amount of ubiquitination by over 2-fold (Fig. 5C). To directly assess the effect of Thr455 phosphorylation on ubiquitination, 293T cells expressing VEC or PIM1 were transfected with HA-HIF-1α WT, T455D, or T455A, treated with MG-132 for 4 hours, and HA-HIF-1α was immunoprecipitated. Overexpression of PIM1 significantly decreased WT HIF-1α ubiquitination compared to controls, whereas PIM1 expression did not alter the ubiquitination of T455D and T455A HA-HIF-1α (Fig. 5D). Moreover, the level of ubiquitination of T455A was similar to that of wild-type HIF-1α, whereas T455D ubiquitination was significantly lower than that of wild-type HIF-1α and similar to the amount of ubiquitination observed for wild-type HIF-1α with PIM1 overexpression (Fig. 5D).

**Figure 5.**
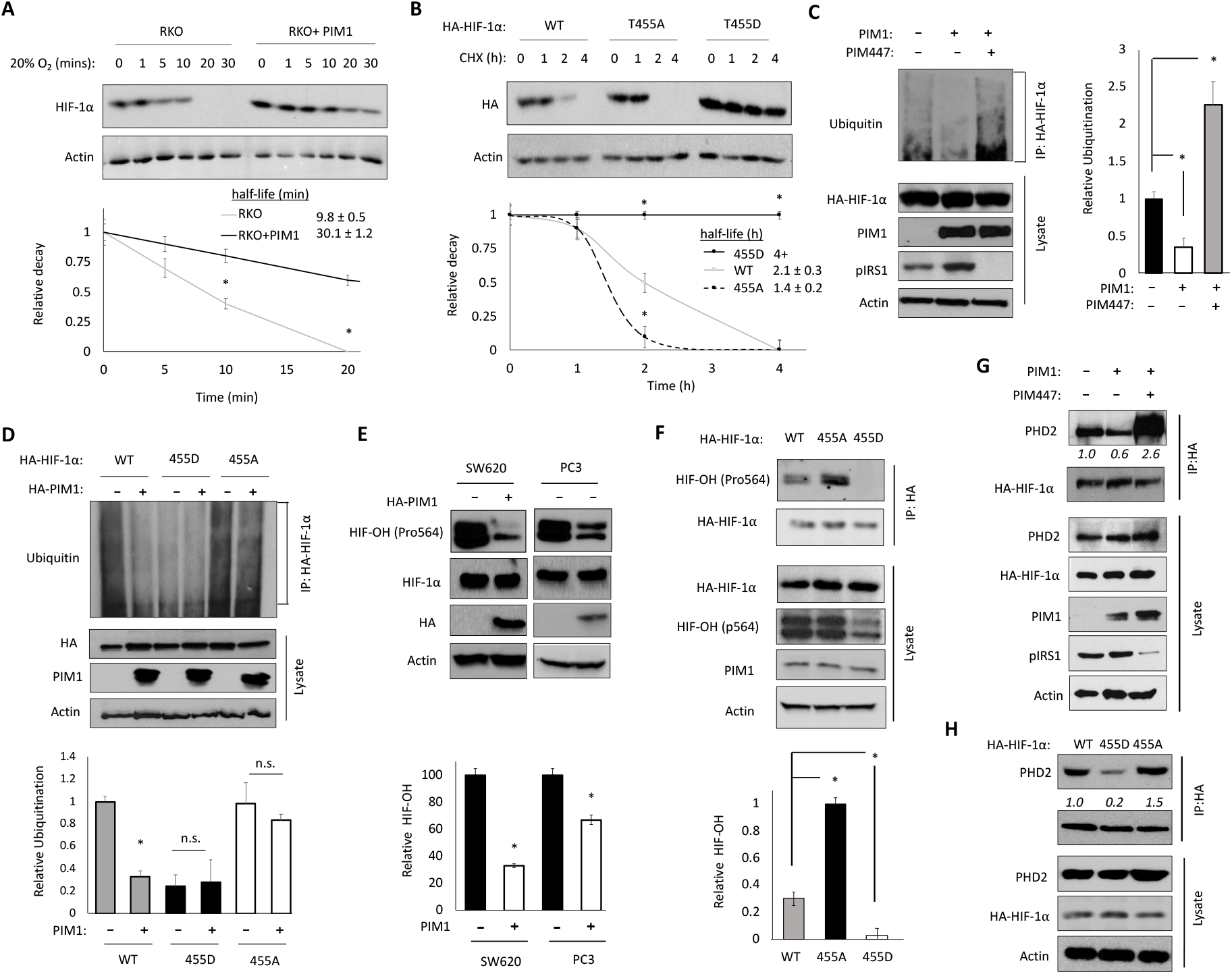
Phosphorylation of HIF-1α at T455 disrupts PHD2 binding and increases HIF-1α stability. A) RKO ± PIM1 were incubated in hypoxia (1.0% O_2_) for 1 h then lysed at different time-points after restoring normal oxygen (20% O_2_). B) 293T cells were transfected with HA-HIF-1α, T455A or T455D and incubated in hypoxia for 4 h prior to treatment with cycloheximide (CHX, 10 μM). Densitometry was used to determine the rate of protein decay. C) 293T ± PIM1 cells were transfected with HA-HIF-1α and treated with MG-132 (10 μM) and DMSO or AZD1208 (3 μM) for 4 h. HIF-1α constructs were immunoprecipitated and ubiquitination was measured by immunoblotting and quantified by densitometry. D) 293T cells ± PIM1 were transfected with HA-HIF-1α, T455D or T455A. E) SW620 and PC3 cells were transfected with HA-PIM1 and lysates were collected. Relative HIF-OH (Pro564) is graphed. F) HA-HIF-1α, T455D or T455A. HA-HIF-1α constructs were immunoprecipitated and blotted for HIF-OH (Pro564). The ratio of hydroxylated to total HIF-1α is graphed. G) 293T ± PIM1 cells were transfected with HA-HIF-1α and treated with DMSO or PIM447 (3 μM) for 4 h. HA-HIF1a constructs were immunoprecipitated and PHD2 was probed by western blotting; relative abundance was calculated by densitometry. H) HA-HIF1a constructs were immunoprecipitated and PHD2 was probed by western blotting; relative abundance quantified below. *p < 0.05, n.s. = not significant, error bars = SEM

Because hydroxylation is the initiating step in the canonical HIF-1α degradation pathway, we next assessed changes in HIF-1α hydroxylation at Pro564. First, SW620 colon and PC3 prostate cancer cells were treated for 4 h with MG132 to allow for the accumulation HIF-1α and hydroxylation was measured by western blotting. Despite there being similar levels of total HIF-1α, hydroxylation of HIF-1α at Pro564 was significantly reduced by PIM1 overexpression (Fig. 5E). In a similar experiment, HA-HIF-1α WT, T455D, or T455A were immunoprecipitated from 293T cells stably expressing VEC or PIM1 (to exclude any endogenous HIF-1α) after 4 h treatment with MG-132 to allow for the accumulation of hydroxylated HIF-1α. The T455D mutant showed no visible hydroxylation, whereas hydroxylation of the T455A mutant was significantly increased compared to WT HIF-1α (Fig. 5F). PHD2 is the primary isoform responsible for hydroxylating HIF-1α. Because Thr455 is located within the ODDD, we hypothesized that phosphorylation at this site may disrupt PHD binding to HIF-1α. To this end, HA-HIF-1α was transfected into 293T cells stably expressing PIM1 or vector control. Cells were treated with MG132 for 4 h, HA-HIF-1α was immunoprecipitated, and PHD2 binding was assessed by immunoblotting. Significantly less PHD2 was bound to HIF-1α in cells overexpressing PIM1 compared to VEC cells (Fig. 5G). To test whether Thr455 phosphorylation alters PHD2 binding, 293T cells were transfected with HA-HIF-1α WT, T455D, or T455A constructs and treated with MG-132 for 4 h prior to immunoprecipitation of HA-HIF-1α variants. Significantly more PHD2 was bound to HIF-1α T455A than wild-type, whereas significantly less was bound to HIF-1α T455D compared to wild-type (Fig. 5H). These data indicate that PIM1-mediated phosphorylation of HIF-1α at Thr455 increases protein stability by blocking PHD binding, hydroxylation, and subsequent proteasomal degradation of HIF-1α.

### The anti-tumor effects of PIM inhibitors depend on downregulation of HIF-1

To confirm the significance of Thr455 phosphorylation *in vivo* and *in vitro*, we next created two homozygous HIF-1α T455D mutant SW620 colon cancer cell lines using CRISPR site-directed mutagenesis. We validated these lines by Sanger sequencing (Fig. S4A). Both of these cell lines showed stable HIF-1α protein in normoxic conditions (Fig. 6A) and had significantly increased expression of HIF-1 target genes (Fig. S4B). To assess cell growth, parental SW620 and HIF-1α T455D mutant cell lines (B24 and C34) were plated at various densities (1000, 2000, and 3000 cells per well) and allowed to grow for 48 h. Then, MTT assays were performed to assess relative cell number. The HIF-1α T455D clones, B24 and C34, displayed significantly increased growth compared to parental SW620 cells (Fig. S4C). Next, cells were treated with increasing doses of AZD1208 and PIM447 for 24 h, and MTT assays were performed to assess cell viability. Parental SW620 cells exhibited a significant and dose-dependent reduction in viability in response to both PIM inhibitors, whereas the B24 and C34 cell lines were less sensitive (Fig. S4D). To confirm that HIF-1α was refractory to PIM inhibition in the T455D mutant cell lines, cells were treated with PIM447 (1 μM) for 4 h. Immunoblotting revealed that PIM447 reduced pIRS1 (S1101), a known PIM substrate, whereas HIF-1α levels were significantly higher and refractory to PIM inhibition in the T455D mutant cell lines (Fig. 6A). To determine whether Thr455 phosphorylation also impacted angiogenesis, CM was harvested from parental and B24 and C34 SW620 cells treated with or without PIM447 for 24 h and used for *in vitro* tube formation assays. CM from both B24 and C34 cells significantly increased both the total tube length and number of branch points compared to parental SW620 CM. Treatment with PIM447 significantly reduced the tube length and number of branch points resulting from SW620 CM compared to DMSO, whereas the mutant cell lines were refractory to the anti-angiogenic effect of PIM inhibitors (Fig. 6B and C). Next, we assessed the tumorigenicity of these cell lines and sensitivity to PIM inhibition *in vivo*. Five million parental SW620, C34, or B24 cells were injected into the flanks of 12 (n=4/group) SCID mice. Tumors were allowed to grow to an average size of 100 mm^3^, and then the mice were randomly segregated into vehicle or AZD1208 (30 mg/kg) treatment groups. Tumor growth was measured by caliper every other day, and tumors were harvested for mRNA and IHC analysis once maximum tumor burden was reached. Both B24 and C34 tumors grew significantly faster than parental SW620 tumors (Fig. 6D and Fig. S5A). Strikingly, treatment with AZD1208 significantly reduced the volume of SW620 tumors but was unable to slow the growth of either the B24 or C34 mutant xenografts (Fig. 6D and Fig. S5A). Hematoxylin and eosin (H&E) staining revealed that AZD1208 disrupted tumor vasculature in the SW620 xenografts, resulting in necrotic tissue, whereas C34 and B24 tumors were highly vascular, regardless of PIM inhibition (Fig. 6E and Fig. S5A and B). Importantly, AZD1208 significantly decreased microvessel density (CD31 staining) in SW620 tumors, whereas no significant decrease in vasculature was observed in mutant tumors after treatment with AZD1208 (Fig. 6F). AZD1208 significantly increased apoptosis (cleaved caspase-3) in SW620, but not in HIF-1α mutant tumors (Fig. 6G). Strikingly, HIF-1α levels were significantly higher in C34 compared to wildtype, and AZD1208 significantly reduced HIF-1α levels in the SW620, but not in the C34 tumors (Fig 6H). RT-PCR analysis of tumor tissue confirmed that PIM inhibition reduced the expression of pro-angiogenic genes *VEGF-A* and *ANGPT4* in SW620 tumors, whereas PIM inhibition had no effect on HIF-1 target genes in B24 and C34 tumors (Fig. 6I). These data indicate that phosphorylation of Thr455 is sufficient to drive angiogenesis and increase tumor growth. Moreover, the fact that a single point mutation in HIF-1α made these tumors largely refractory to PIM inhibition suggests that the anti-tumor effects of small molecule PIM inhibitors is primarily due to their ability to downregulate HIF-1 and reduce angiogenesis.

**Figure 6.**
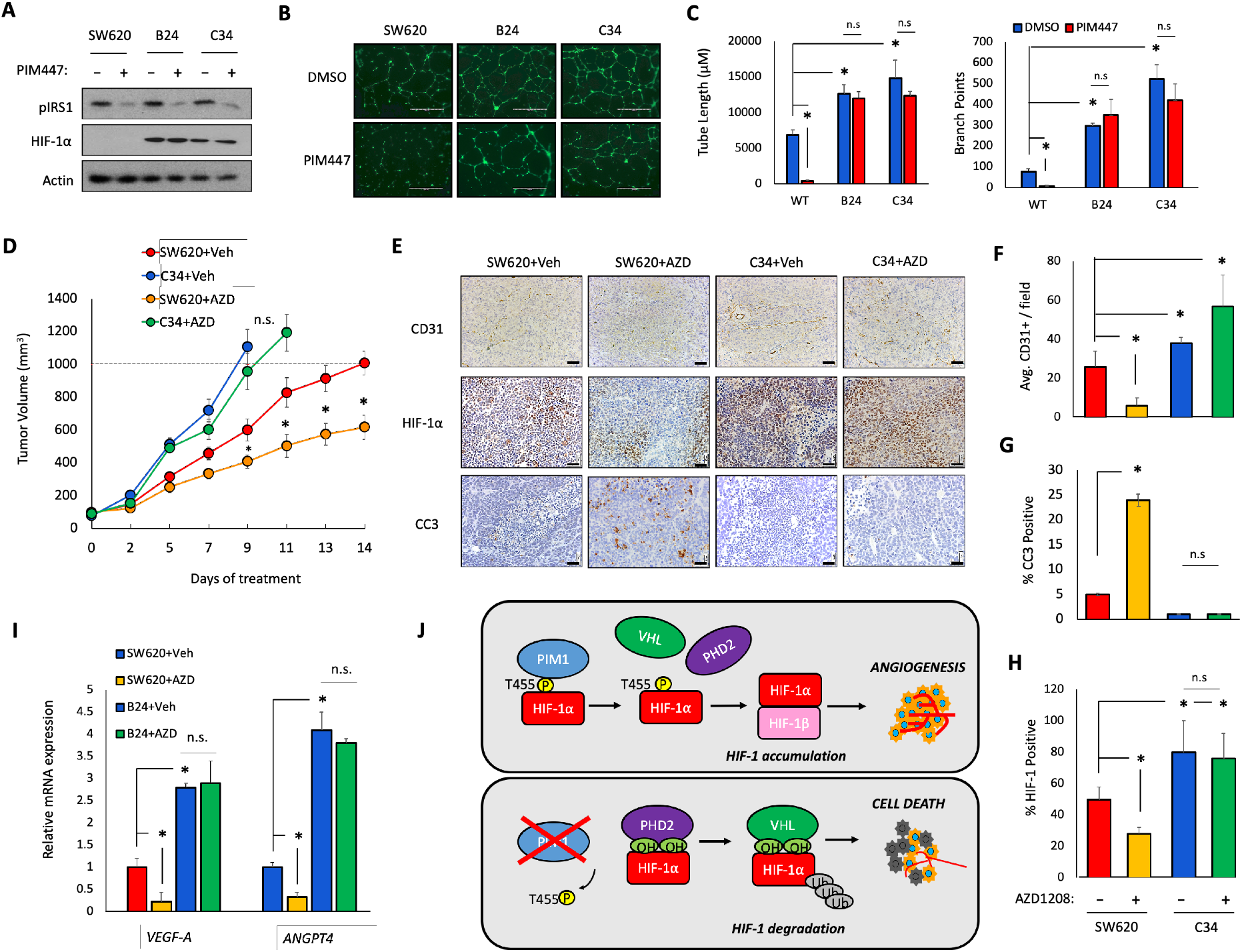
HIF-1α-T455D CRISPR mutants increase tumor growth and are resistant to PIM inhibition. Two homozygous HIF-1α-T455D SW620 cell lines were generated via CRISPR site directed mutagenesis (B24 and C34). A) SW620, B24, and C34 were treated with PIM447 (3 μM) for 6 h. B) Representative images of tube formation after 24 h incubation in the indicated CM. C) Mean tube length and F) total branch points were quantified. D) Mice (n=4/group) were injected with 5×10^6^ SW620 or C34 cells, treated with vehicle or AZD1208 (30 mg/kg), and tumor volume was measured over time. E-H) IHC staining and quantification of the indicated markers in tumors from each cohort. I) RNA was harvested from tumor tissue and mRNA expression was measured by qRT-PCR. J) Model depicting the mechanism and physiological outcome of PIM1-mediated phosphorylation of HIF-1α. *p < 0.05, Scale bar = 50 μm, n.s. = not significant, error bars = SEM.

## Discussion

Angiogenesis is a hallmark of cancer that is required to sustain tumor survival and growth (Hanahan and Weinberg, 2011). HIF-1 activation is arguably the most important driver of angiogenesis in solid tumors. In clinical settings, immunohistochemistry analysis of patient’s biopsy specimens exhibited dramatic HIF-1 overexpression in common human cancers, which leads to increased metastasis and mortality rates (Baba et al., 2010; Lee et al., 2007; Liu et al., 2015; Tong et al., 2016). Here, we are the first to demonstrate that PIM1 is sufficient to activate HIF-1 in normoxic conditions and promote angiogenesis (Fig. 2), establishing a new mechanism by which PIM kinases promote tumor progression, independent of their established roles in regulating cell cycle progression and survival. Therefore, overexpression of PIM1 provides a novel explanation for the constitutive activation of HIF-1 in tumors that is frequently observed in the absence of hypoxia. We identified a novel post-translational modification that controls the proteasomal degradation of HIF-1α (Fig. 3 and 4). Notably, the sequence surrounding HIF-1α at Thr455 does not closely resemble the phosphorylation consensus motif predicted for PIM kinases. Several established PIM1 substrates also have atypical phosphorylation sites, making it unclear of how complete the described consensus sequences are for predicting PIM substrates. Mechanistically, phosphorylation of HIF-1α by PIM1 at Thr455 prevents the binding of PHDs and subsequent hydroxylation of HIF-1α, ultimately reducing its ubiquitination by VHL and degradation by the 26S proteasome (Fig. 6L). HIF-1α is phosphorylated at Thr455 in human cancer cell lines and this site is sensitive to PIM1 activity but may be mor widely regulated. For example, earlier work in pancreatic cancer showed that PIM3 correlated with increased VEGF expression, suggesting that other PIM isoforms may also be able phosphorylate HIF-1α at Thr455 (Wang et al., 2013). Moreover, protein kinase A has been reported to phosphorylate HIF-1α at Thr455, although this site was not characterized (Bullen et al., 2016). Additionally, another proteomic-based screen searching for targets of aurora kinase inhibitors found significantly decreased phospho-Thr455 after treatment with aurora kinase inhibitors (AZD1152 and ZM447439) (Kettenbach et al., 2011). Therefore, phosphorylation of Thr455 appears to be a widely used mechanism through which multiple kinases regulate HIF-1α protein levels and HIF-1 activation. Unfortunately, HIF-1α itself has yet to be crystallized thus we are unable to clearly speculate the consequences of T455 phosphorylation to HIF structure. Additionally, T455 is not a canonical PIM site (RXRHXS), but this, again, is difficult to study without HIF protein folding information. As well, other sites have recently been identified as PIM phospho-sites despite no canonical sequence being present, including, but not limited to, PIM1 phosphorylating itself (Bullock et al., 2005). Future studies warrant further understanding of how PIM kinases recognize alternative substrate sequences and what determines the specificity of these targets.

PIM1 is elevated in many solid tumors and feeds into several pro-tumorigenic signaling pathways (Brault et al., 2010; Rebello et al., 2018; Warfel and Kraft, 2015). Here, we establish that heightened expression of PIM1 was sufficient to initiate and sustain angiogenesis in solid tumors (Fig. 1 and 2). Overexpression of PIM1 in multiple xenograft tumor models increased tumor volume, as well as microvessel density (CD31) and vascular perfusion (MRI). Importantly, analysis of prostate cancer TMAs revealed a significant correlation between PIM1 and microvessel density, confirming that our *in vitro* and *in vivo* data translate to human cancers. Tumor vasculature is also the primary route for tumor cell dissemination (Bielenberg and Zetter, 2015; Tannock, 1968). Several preclinical and clinical studies in various solid tumor models show that upregulation of PIM isoforms is associated with increased metastasis and PIM inhibitors alone prevent metastasis (Casillas et al., 2018; Qu et al., 2016; Santio et al., 2015). Specifically, a recent clinical study revealed that PIM1 is overexpressed at a high frequency in circulating tumor cells from patients with castrate-resistant prostate cancer (Markou et al., 2020). Our results suggest that studies that the pro-metastatic properties of PIM could be linked to increased tumor vascularization. In contrast to control tumors, overexpression of PIM1 in xenograft tumors lacking HIF-1α did not increase tumor growth, proliferation, or angiogenesis, indicating that the pro-tumorigenic effects of PIM1 are largely dependent on stabilization of HIF-1α and heightened angiogenesis (Fig 2). In addition, CRISPR cell lines containing a phospho-mimetic at Thr455 were completely refractory to the anti-tumor effects of PIM inhibitors *in vivo*. AZD1208 significantly reduced tumor volume, induced cell death, and reduced angiogenesis in parental xenografts but was unable to induce any of these physiological consequences in the T455D xenografts (Fig 6). Thus, the anti-tumor effect of PIM inhibitors is dependent on their ability to downregulate HIF-1 and reduce angiogenesis (Fig. 6). Synergistic anti-tumor effects have been reported when combining PIM inhibitors with a plethora of targeted and chemo-therapies, regardless of their mechanism of action (Casillas et al., 2018; Cen et al., 2014; Hammerman et al., 2005; Kurmasheva et al., 2006; Moody et al., 2015). Thus, our results suggest that the wide-ranging efficacy observed when combining PIM inhibitors and cytotoxic therapies can be largely attributed to their anti-angiogenic effects. We previously reported that PIM inhibitors selectively kill hypoxic tumor cells by increasing cellular reactive oxygen species in a HIF-1-independent manner (Warfel et al., 2016a). This research expands our working model to include HIF-1 in the *in vivo* setting. PIM inhibitors reduce angiogenesis by inhibiting HIF-1, which exacerbates the effects of hypoxia and renders tumor cells more susceptible to their cytotoxic effects (Fig. 6J).

Successful therapeutic targeting of transcription factors has proven challenging. However, reducing the expression of key transcription factors through post-translational modifications represents a promising approach (Bushweller, 2019; Filtz et al., 2014; Huang et al., 2013). Ongoing efforts to block angiogenesis as a treatment strategy have been fraught with disappointment, in many cases due to an increase in hypoxia associated with more aggressive disease. While many small molecules have been reported as HIF-1α inhibitors, no clinically approved selective HIF-1α inhibitor has been reported to date. Those that have been tested in the clinic has shown modest benefit (Fallah and Rini, 2019), suggesting that alternative, HIF-independent pro-survival pathways are able to compensate or overcome HIF-1 inhibition. Taken in the context of our and other research studies, PIM is poised to fill this role. Blocking PIM represents a promising approach to overcome hypoxia-mediated therapeutic resistance due to both HIF-dependent and HIF-independent effects. Taken together with our previous studies showing that co-inhibition of PIM1 and VEGF is a viable approach to treating solid tumors (Casillas et al., 2018; Chauhan and Warfel, 2018), this work provides further rationale for how this strategy works at the molecular level and identifies HIF-1α phosphorylation at Thr455 as a potential biomarker for the efficacy of PIM inhibitors in human tumors. Therefore, further translation of PIM inhibitors in combination with approved therapies is warranted in solid tumors as a new strategy to inhibit HIF-1 and angiogenesis while simultaneously targeting the hypoxic tumor cell population that is commonly associated with acquired resistance to therapy and treatment failure.

## Acknowledgments

We would like to thank the core services of In vivo Imaging, EMSR, TACMASR, Gene Editing, Mass Spectrometry, and the Biostatistics Shared Resources at The University of Arizona for their help and support. We would also like to thank Dr. Dan Buster for performing *in vitro* kinase assays and Brenda Baggett and Dr. Marty Pagel for assisting with *in vivo* imaging. Studies were supported by funding from the National Cancer Institute (T32CA009213) on behalf of A.L.C., American Cancer Society (RSG-16-159-01-CDD) to N.A.W., and Department of Defense PCRP (W81XWH-19-1-0455) to N.A.W. Cancer Center Support grant from the National Institute of Health (P30CA023074) also supported this research.

## Author Contributions

A.L.C and N.A.W. conceived the project and developed experimental methodology. A.L.C led the study under the supervision of N.A.W., A.E.C., C.K.M., A.L.C., S.C.C., A.G.S, R.K.T., A.N.C., C.J.J., P.R.L., and N.A.W. performed experiments and contributed to data analysis. A.L.C., P.R.L., and N.A.W. created figures and wrote the manuscript with input from the other authors.

## STAR Methods

### Key resources table

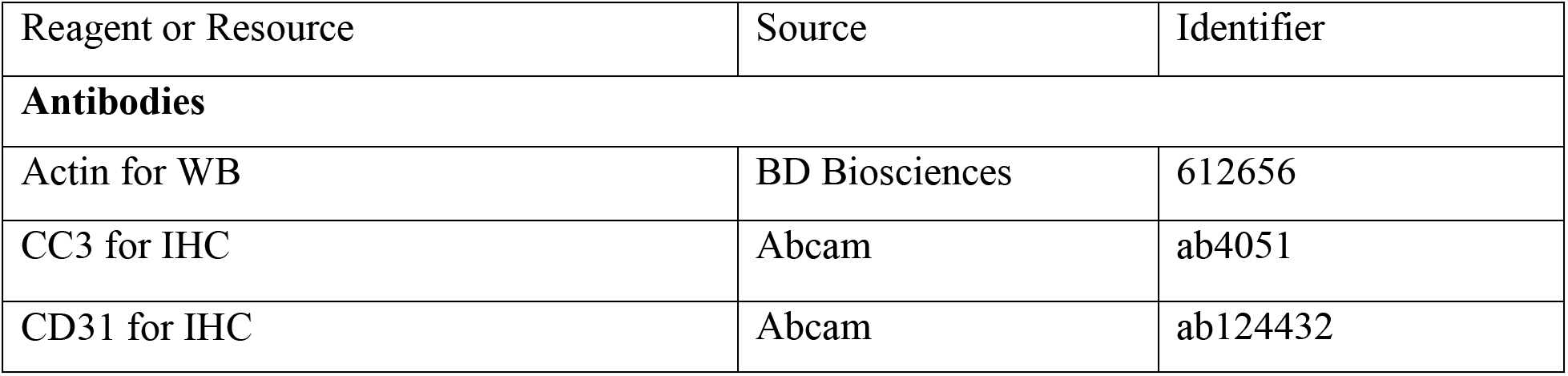
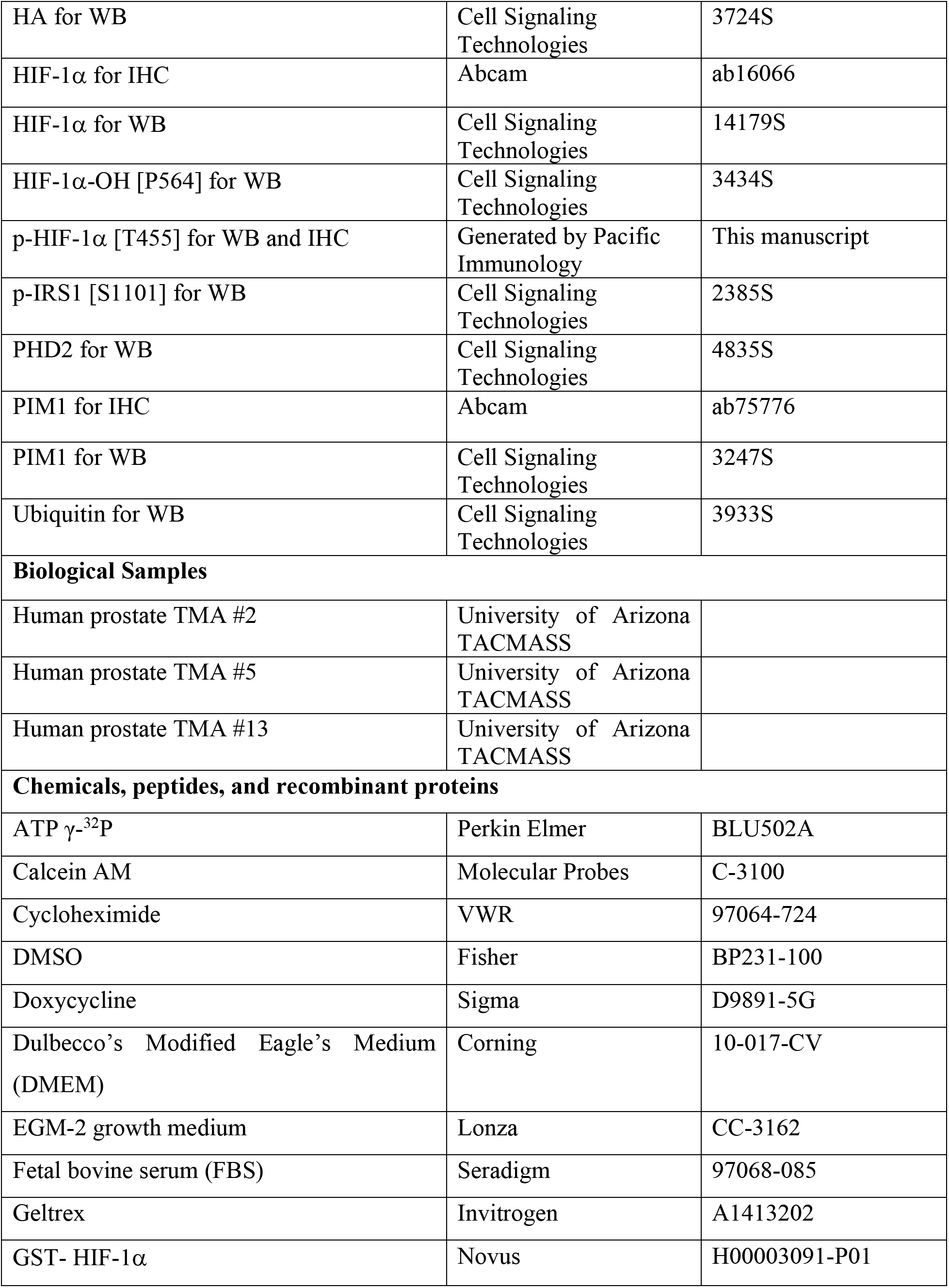
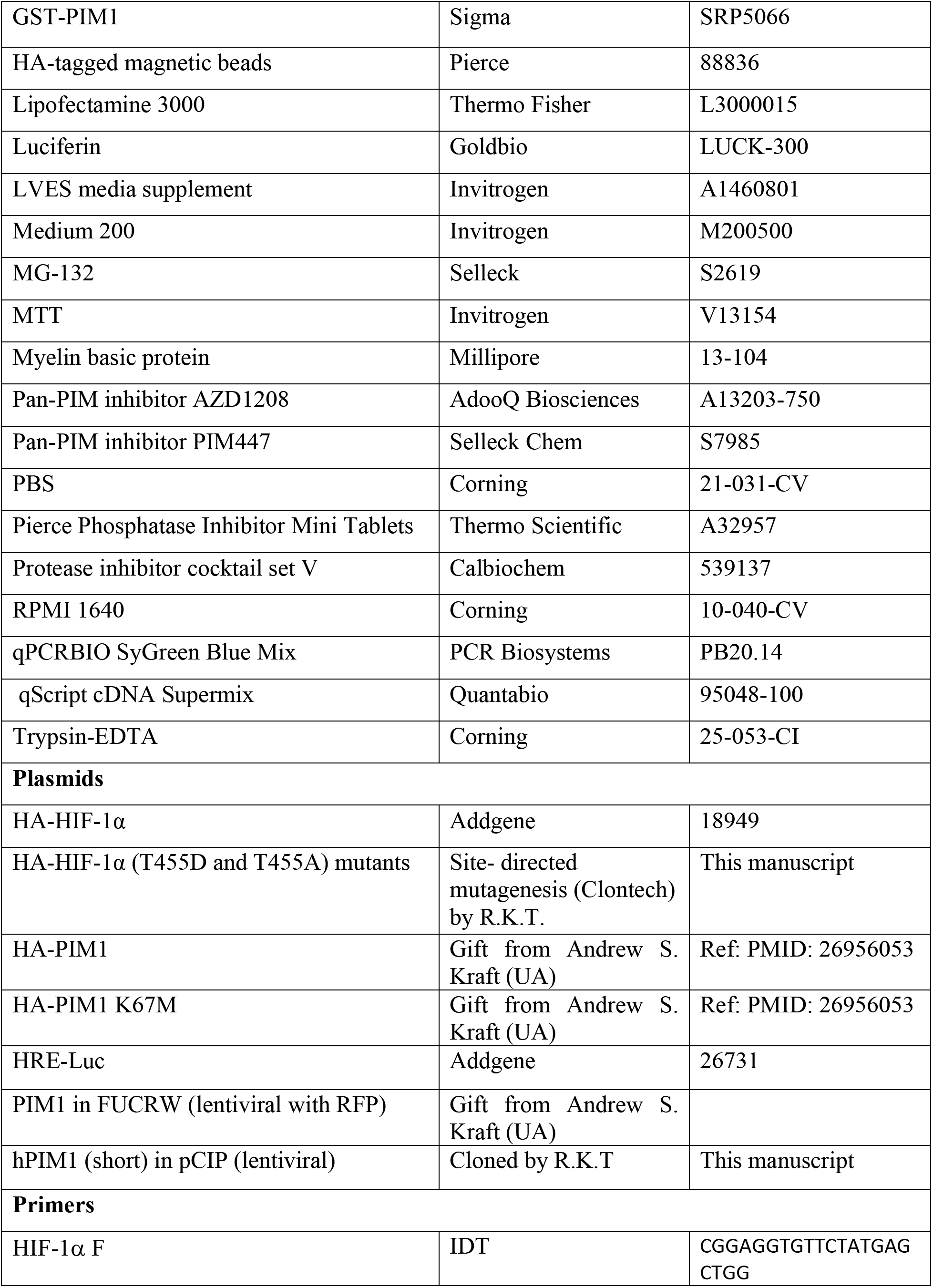
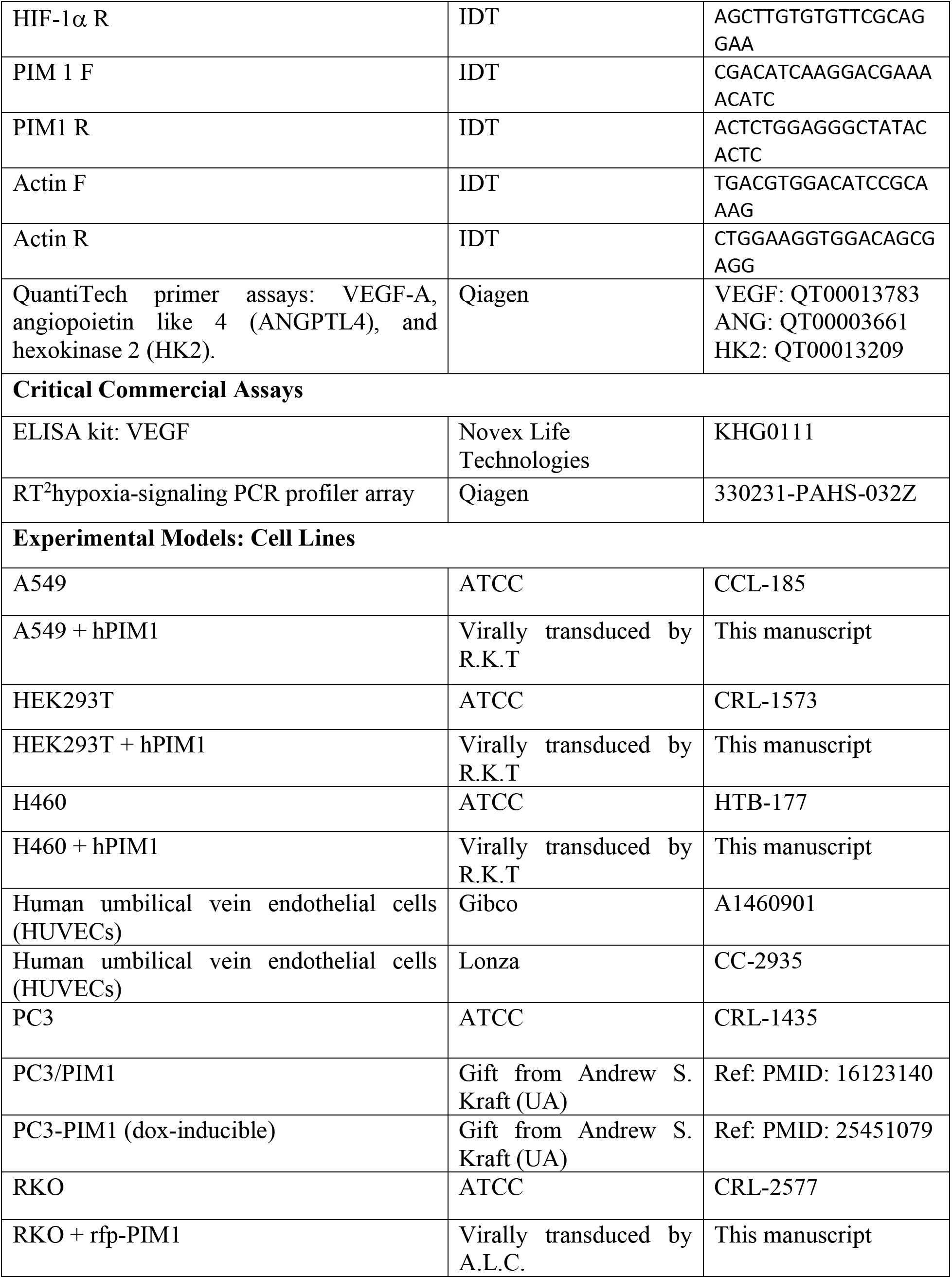
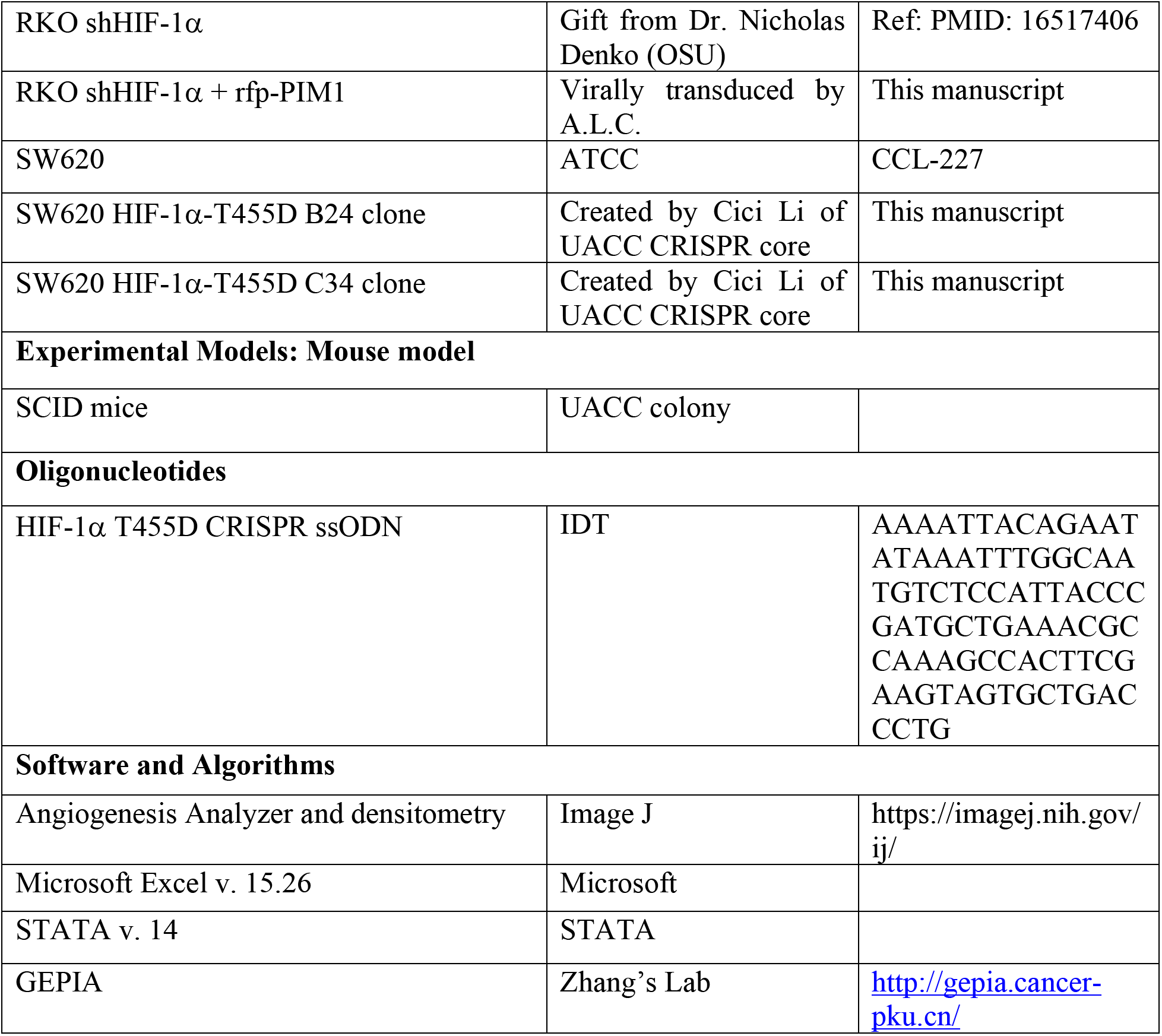

### Lead contact and Materials Availability

Further information and request for resources and reagents should be directed to and will be fulfilled by the Lead Contact, Noel A. Warfel.

## Cell lines

Parental and genetically modified A549, H460, HEK293T, RKO, and SW620 cells were maintained in DMEM medium containing 10% FBS. SW620 HIF-1α-T455D clone B24 and clone C34 cell lines were generated by CRISPR-cas9 mediated mutagenesis using ssODN AAAATTACAGAATATAAATTTGGCAATGTCTCCATTACCCGATGCTGAAACGCCAAAGCCACTTCGAAGTAGTGCTGACCCTG and CRISPR sgRNA5’GGCTTTGGCGTTTCAGCGGT – 3’. PC3/VEC and PC3/PIM1 cell lines were maintained in RPMI medium containing 10% FBS. Human umbilical vein endothelial cells (HUVECs) from Gibco were cultured in complete Med-200 containing 1X LVES media supplement (Invitrogen), while HUVEC cells from Lonza were cultured in EGM-2 medium with kit supplements added (Lonza). All cells were cultured at 37°C in 5% CO_2_, routinely screened for mycoplasma, and authenticated by short tandem repeat DNA profiling performed by the University of Arizona Genetics Core Facility and were used for fewer than 50 passages. When appropriate, cells lines were cultured in a hypoxic environment (1% O_2_, 5% CO_2_, 94% N_2_) using an InVivO_2_ 400 hypoxia workstation (Baker Ruskinn).

## Antibodies

The following western blot antibodies were used: Actin (BD Biosciences); and HA, HIF-1α, HIF-1α-OH [P564], p-IRS1 [S1101], PHD2, PIM1 and Ubiquitin (Cell Signaling Technologies). The p-HIF-1α [T455] antibody was generated by Pacific Immunology for western blot and IHC in this manuscript. The following antibodies were also used in IHC: CC3, CD31, HIF-1α and PIM1 (ab245417)(Abcam).

## Kinase assays

### In vitro

Recombinant full-length GST-PIM1 was incubated with recombinant GST-HIF-1α, GST (negative control), or dephosphorylated myelin basic protein (MBP) in the presence or absence of AZD1208 (100 nM) and ^32^P-labeled ATP in kinase assay buffer (20 mM MOPS [pH 7.0] containing 100 mM NaCl, 10 mmol/L MgCl_2_, and 2 mmol/L dithiothreitol). Reactions were incubated at 37°C for 60 min prior to the addition of 5× Laemmli sample buffer. Samples were separated by SDS-PAGE and gels were stained with Coomassie prior to autoradiography or preparation for mass spectrometry.

### In vivo

HA-HIF-1α or its mutant (T455A) were immunoprecipitated with anti-HA antibodies. Immune complexes were washed three times in lysis buffer, then washed twice in 1 × kinase buffer and incubated with 0.1 μg of recombinant active PIM1 and 100 μmol/L of ATP for 30 min at 25°C. Reactions were stopped by washing twice in a cold kinase buffer and boiling in 2 × SDS loading buffer. The sample was separated on an SDS-polyacrylamide gel and subjected to Western blot analysis with anti-phospho HIF-1α (T455) antibody.

## DCE-MRI studies

Two million PC3-VEC or PC3-PIM1 cells were injected subcutaneously into the rear flanks of 8 mice (n=4/group) and tumor volume was measured by caliper. Tumors were allowed to grow to ~300 mm^3^ before initiating MRI studies using a 7T Bruker Biospec MRI instrument. Prior to the MRI scan, each mouse had a 27G catheter placed in the tail vein and was anesthetized with 1.5-2.5% isoflurane in O_2_ carrier gas. Physiologic respiration rate and core body temperature were monitored throughout the MRI session. All animals were imaged while maintaining their temperature at 37.0 ± 0.2 °C using warm air controlled by a temperature feedback system (SA Instruments). The T1 relaxation time of each tissue of interest (30) was measured by acquiring a series of spin-echo MR images, variable TR and a RARE protocol using the following parameters: TR = 150, 300, 350, 500, 700, 900, 1200, 2000, 3000 and 6000 msec; TE = 9.07 msec; NEX = 1; RARE factor = 2; slice thickness = 1.0 mm; FOV = 2.0 cm2; linear encoding order; matrix = 128 x 128; in-plane spatial resolution = 0.23 mm2; hermite excitation pulse = 90° for 2.7 msec duration with 2700 Hz bandwidth; and hermite refocusing pulse = 180° for 1.71 msec duration with 2000 Hz bandwidth. The T1 time for each image voxel was estimated by non-linear regression of the variable TR signal to the following equation: (1) MZ (t) = MZ (1 – e − TR/T1). A series of dynamic images were acquired using a Spoiled Gradient-echo MRI protocol with the following parameters: TR= 50 msec; TE= 8.07 msec; NEX = 1; excitation pulse = 158.9°; FOV = 2.0 cm2; in-plane spatial resolution = .23 mm2; matrix = 128 x 128; slice thickness = 1.0 mm, for a single slice centered in the tumor. Each image set was acquired in 6.4 sec and repeated 150 times for a total acquisition time of 16 min. An initial set of baseline images were acquired for 30 sec. prior to I.V. injection of 50 μL of MultiHance (Bracco Diagnostics Inc.) over a minute, which corresponds to a dose of 0.40 mMKg-1 for a 20 g mouse. All images were analyzed using the linear reference region model (Eq. [1]) after transforming the MRI signals to concentrations. Muscle in the left thigh of each mouse was used as the reference region. Upon sacrifice, tissue was harvested and stained by IHC for PIM1 and CD31.

(1) ΔR1,TOI (T) = RK_trans_ ⋅ ΔR1,RR(T) + Ktrans,TOI/e,RR⋅ 0∫T ΔR1,RR(t) dt − kep,TOI ⋅ 0∫T ΔR1,TOI (t) dt

## *In vivo* studies

### Fig. 2

Five million parental or HIF knockdown or PIM1 overexpressing or both RKO cells in PBS were injected subcutaneously into the rear flanks of twelve mice. Tumor volume was measured overtime by caliper. At day 20, mice were injected with 2 nmol of Angiosense 750EX via tail vein injection. Twenty-four hours later, mice were anesthetized and imaged in a Lago Bioluminescence imager (Spectral Instruments). Mice were sacrificed when tumor burden (1000mm^3^) or day 24 of the study. Tumors were harvested for mRNA analysis and IHC staining with HIF-1α, CC3, and CD3.

### Fig. 6

Five million parental, B24, or C34 SW620 cells in PBS were injected subcutaneously into the rear flanks of twelve mice. Once the tumors reached a volume of approximately 100 mm^3^, the mice were randomized for daily treatment with vehicle or AZD1208 (30 mg/kg/day, p.o.) for up to 2 weeks or until tumor burden reached a maximum volume of 2 cm^3^. Tumors were measured every 2-3 days via caliper. Upon sacrifice, tumors were harvested for IHC staining with PIM1, HIF-1α, and CC3, and mRNA was collected for analysis.

For both, the following formula was used to calculate tumor volume by caliper measurements: V = (tumor width)^2^ × tumor length/2.

## In vitro angiogenesis assay

Ten thousand HUVEC cells were plated onto 50 μL Geltrex matrix (Invitrogen) with 100 μL CM. CM from cancer cells was collected by adding 2 mL of DMEM or RPMI + 0.5% FBS to 50,000 cells for 48 h before collection. Tubes were allowed to form for up to 6 h (Lonza kit) or 24 hours (Gibco kit) and then were stained with calcein AM and analysis was performed using ImageJ.

## Protein extraction and immunoblotting

Cultured cells were trypsinized and lysed in NP-40 lysis buffer (150 mM sodium chloride, 1% NP-40, 50 mM Tris pH 8.) with protease and phosphatase inhibitors. Proteins were separated via SDS-PAGE, transferred to PVDF membrane, and probed using the indicated antibodies.

## ELISA

CM from cancer cells was collected by adding 2 mL of DMEM + 0.5% FBS to 50,000 cells for 48 hours before collection. Media was spun down to pellet cell debris and sterile-filtered before enzyme-linked immunosorbent assay (ELISA). ELISA was performed according to manufacturer’s procedure (Novex by Life Technologies) and plates were read at 450 nm.

## MTT

### Growth

Ten, twenty and thirty thousand cells were plated onto a 96-well plate and allowed to grow for 48 h before MTT assay (Invitrogen).

### Response to drug

Twenty thousand cells were plated onto a 96-well plate and allowed to grow for 24 h prior to addition of the drug, and MTT assays were performed after 24 h incubation with the indicated drugs. For MTT assays, media was removed from the cells by aspiration and 50 μL serum-free DMEM and 50 μL MTT solution were added to each well, and plates were incubated for 4 h at 37°C. After incubation, 150 μL of DMSO was added to each well for 15 minutes and absorbance was read at 540 nM.

## qRT-PCR

Hypoxia-responsive gene expression was measured and quantified using RT^2^ hypoxia-signaling PCR profiler arrays using the manufacturer’s software (Qiagen). All other qRT-PCR reactions were performed using qPCRBIO SyGreen Blue Mix (PCR Biosystems), according to the manufacturer’s protocol. Validated primer sets (QuantiTech primer assays; Qiagen) for each of the following genes were purchased to measure gene expression: *VEGF-A*, angiopoietin like 4 (*ANGPTL4*), and hexokinase 2 (*HK2*). HIF-1α, PIM1 and Actin primers were purchased from IDT. Actin was used to normalize.

## Mass spectrometry

### In-gel digestion

Gel slices were in-gel digested with trypsin (Pierce Biotechnology, Rockford, IL) for 3 h at 37°C using ProteaseMax™ Surfactant trypsin enhancer following reduction and alkylation with dithiothreitol and iodoacetamide, respectively, according to the manufacturer’s instructions (Promega Corporation, Madison, WI).

### Mass spectrometry and database search

were performed as previously described (Downs et al., 2018). Briefly, LC-MS/MS analysis was carried out using a Q Exactive Plus mass spectrometer (Thermo Fisher Scientific) equipped with a nanoESI source. Peptides were eluted from an Acclaim Pepmap™ 100 precolumn (100-μm ID × 2 cm, Thermo Fischer Scientific) onto an Acclaim PepMap™ RSLC analytical column (75-μm ID × 15 cm, Thermo Fischer Scientific) using a 5% hold of solvent B (acetonitrile, 0.1% formic acid) for 15 minutes, followed by a 5–22% gradient of solvent B over 105 minutes, 22–32% solvent B over 15 minutes, 32–95% of solvent B over 10 minutes, 95% hold of solvent B for 10 minutes, and finally a return to 5% of solvent B in 0.1 minutes and another 14.9 minute hold of solvent B. All flow rates were 300 nL/min using a Dionex Ultimate 3000 RSLCnano System (Thermo Fischer Scientific). Solvent A consisted of water and 0.1% formic acid. Data dependent scanning was performed by the Xcalibur v 4.0.27.19 software using a survey scan at 70,000 resolution scanning mass/charge (m/z) 400–1600 at an automatic gain control (AGC) target of 3e6 and a maximum injection time (IT) of 100 msec, followed by higher-energy collisional dissociation (HCD) tandem mass spectrometry (MS/MS) at 27nce (normalized collision energy), of the 10 most intense ions at a resolution of 17,500, an isolation width of 1.5 m/z, an AGC of 2e5 and a maximum IT of 50 msec. Dynamic exclusion was set to place any selected m/z on an exclusion list for 20 seconds after a single MS/MS. Ions of charge state +1, 7, 8, >8 and unassigned were excluded from MS/MS, as were isotopes. Tandem mass spectra were extracted from Xcalibur ‘RAW’ files and charge states were assigned using the ProteoWizard 3.0 msConvert script using the default parameters. The fragment mass spectra were then searched against the human SwissProt_2018 database (20413 entries) using Mascot (Matrix Science, London, UK; version 2.6.0) using the default probability cut-off score. The search variables that were used were: 10 ppm mass tolerance for precursor ion masses and 0.5 Da for product ion masses; digestion with trypsin; a maximum of two missed tryptic cleavages; variable modifications of oxidation of methionine and phosphorylation of serine, threonine, and tyrosine. Cross-correlation of Mascot search results with X! Tandem was accomplished with Scaffold (version Scaffold_4.8.2; Proteome Software, Portland, OR, USA). Probability assessment of peptide assignments and protein identifications were made through the use of Scaffold. Only peptides with ≥ 95% probability were considered.

## Quantification and Statistical Analysis

All *in vitro* experiments, including *in vitro* angiogenesis assays, western blots, MTT assays and RT-PCR analysis were conducted 3 independent experiments with at least 3 replicates each. Two-way t-test and linear regression analysis by Pearson correlation were used to compare differences between two groups. Two-way ANOVA was used to analyze differences among groups with multiple independent variables. Tumor growth was analyzed by fitting a mixed linear model of tumor volume vs. time for each mouse. All data are presented as the mean ± SE, and a p < 0.05 was considered to be statistically significant. Microsoft Excel and STATA15 were used for analyses.

## Supplemental Data

**Supplemental Figure 1.**
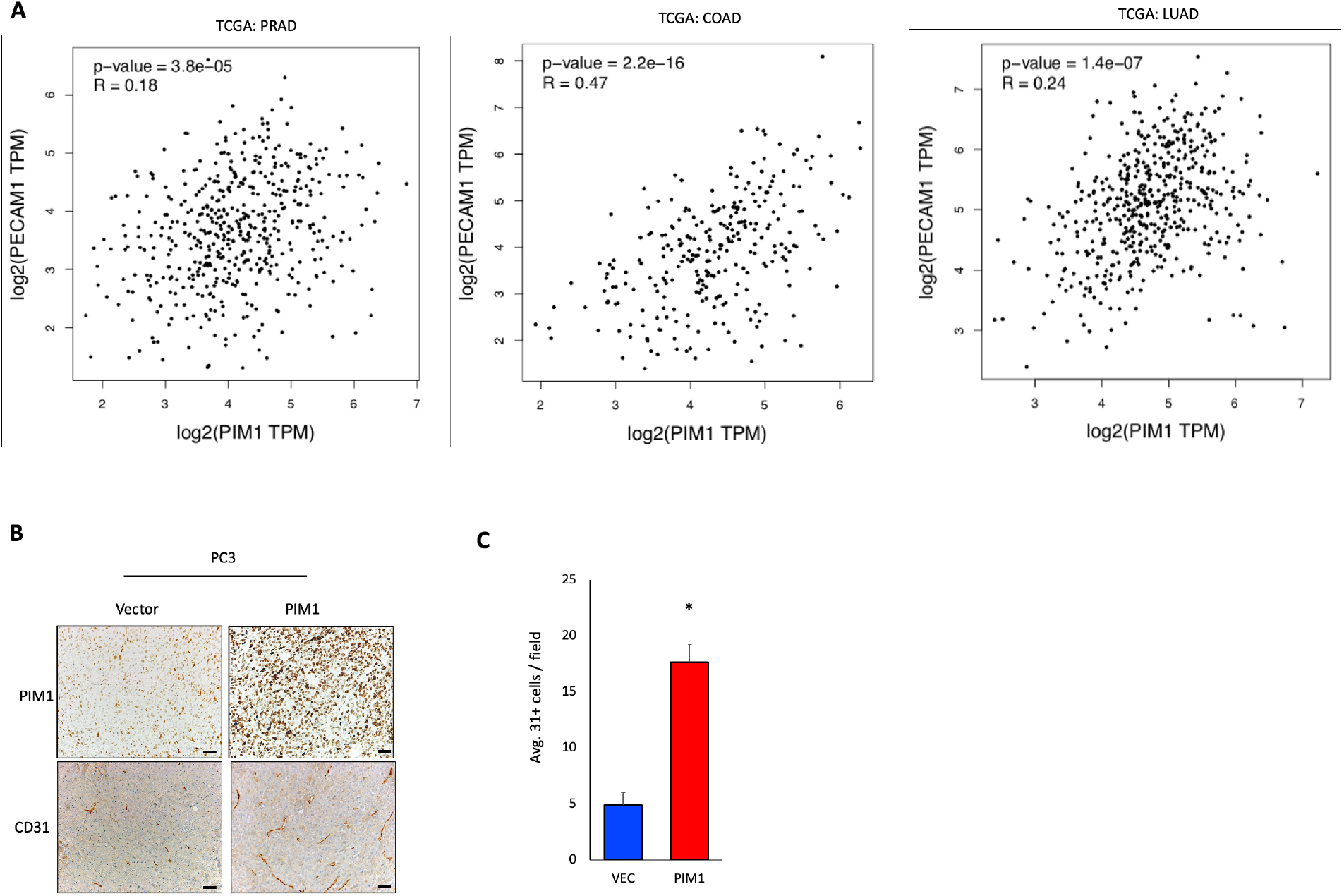
PIM1 is correlated with angiogenesis in cancer. A) Association of PECAM1 and PIM1 transcript levels by Pearson correlation in TCGA prostate (PRAD), colon (COAD), and lung (LUAD) adenocarcinoma datasets. B) Immunostaining of PC3/VEC or PC3/PIM1 tumors C) Average CD31+ vessels calculated from multiple fields. *p < 0.05, error bars = SEM.

**Supplemental Figure 2.**
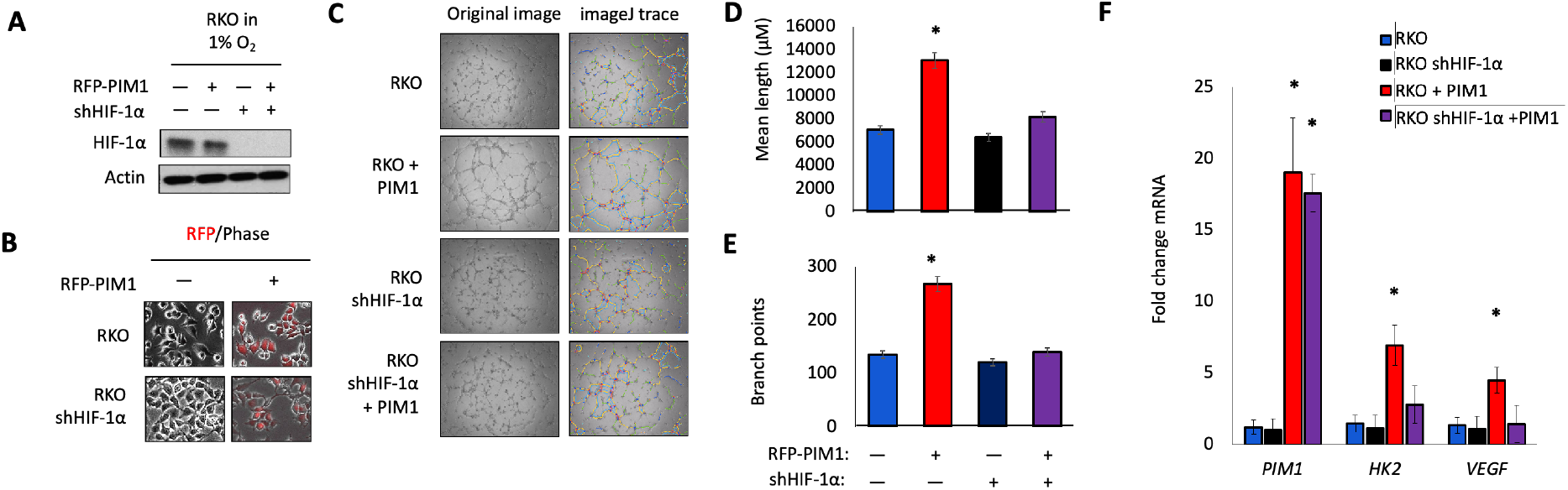
PIM promotes angiogenesis in a HIF-1 dependent manner. A) RKO and RKO shHIF-1α cells were cultured in hypoxia for 4 h and lysates were collected for immunoblotting. B) Representative DIC and RFP images of RKO and RKO shHIF-1α after transduction with RFP-PIM1. C) Tube formation assays using the indicated conditioned media. Representative images and imageJ traces used for quantification of D) mean tube length and E) number of branch points. H) RNA was harvested from tumor tissue and the expression of HIF-1 target genes were quantified by qRT-PCR. *p < 0.05, error bars = SEM.

**Supplemental Figure 3.**
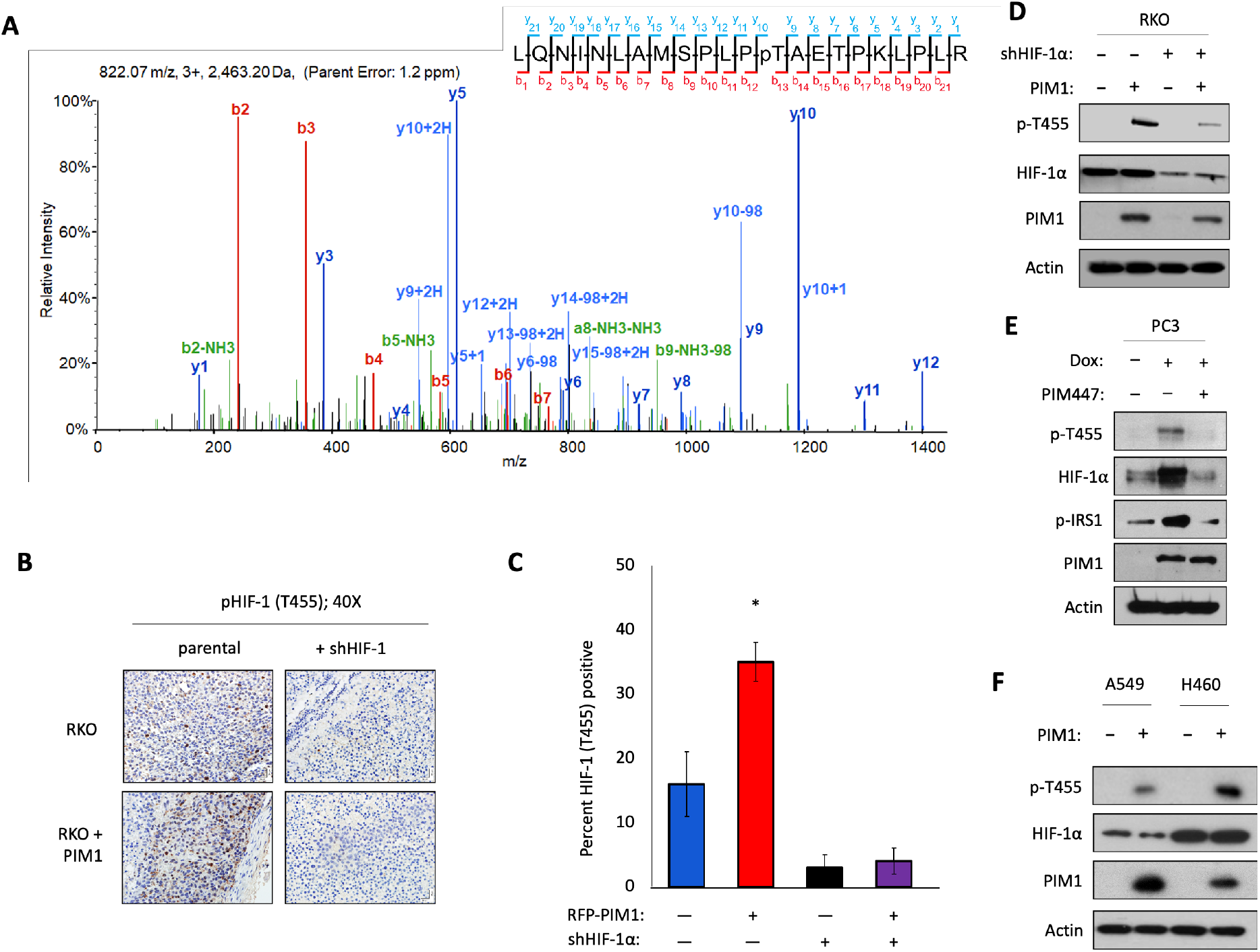
PIM1 phosphorylates HIF-1α at Thr455 *in vivo*. A) Spectra from mass spectrometry analysis of HIF-1α from cells overexpressing PIM1 displays phosphorylation of HIF-1α at Thr455. B) Representative images of p-HIF-1α (T455) immunostaining of tumors from Fig 2H with) C) Quantification of IHC. D) The indicated RKO cell lines were treated with MG-132 (10 μM) for 4 h. E) Dox-PIM1 PC3 cells were treated for 24 h with Dox and pretreated with MG-132 for 4 h followed by DMSO or PIM447 (1 μM) for 6 h; and E) A549 and H460 lung cancer cells ± PIM1 were treated with MG132 for 3 h. *p < 0.05, error bars = SEM.

**Supplemental Figure 4.**
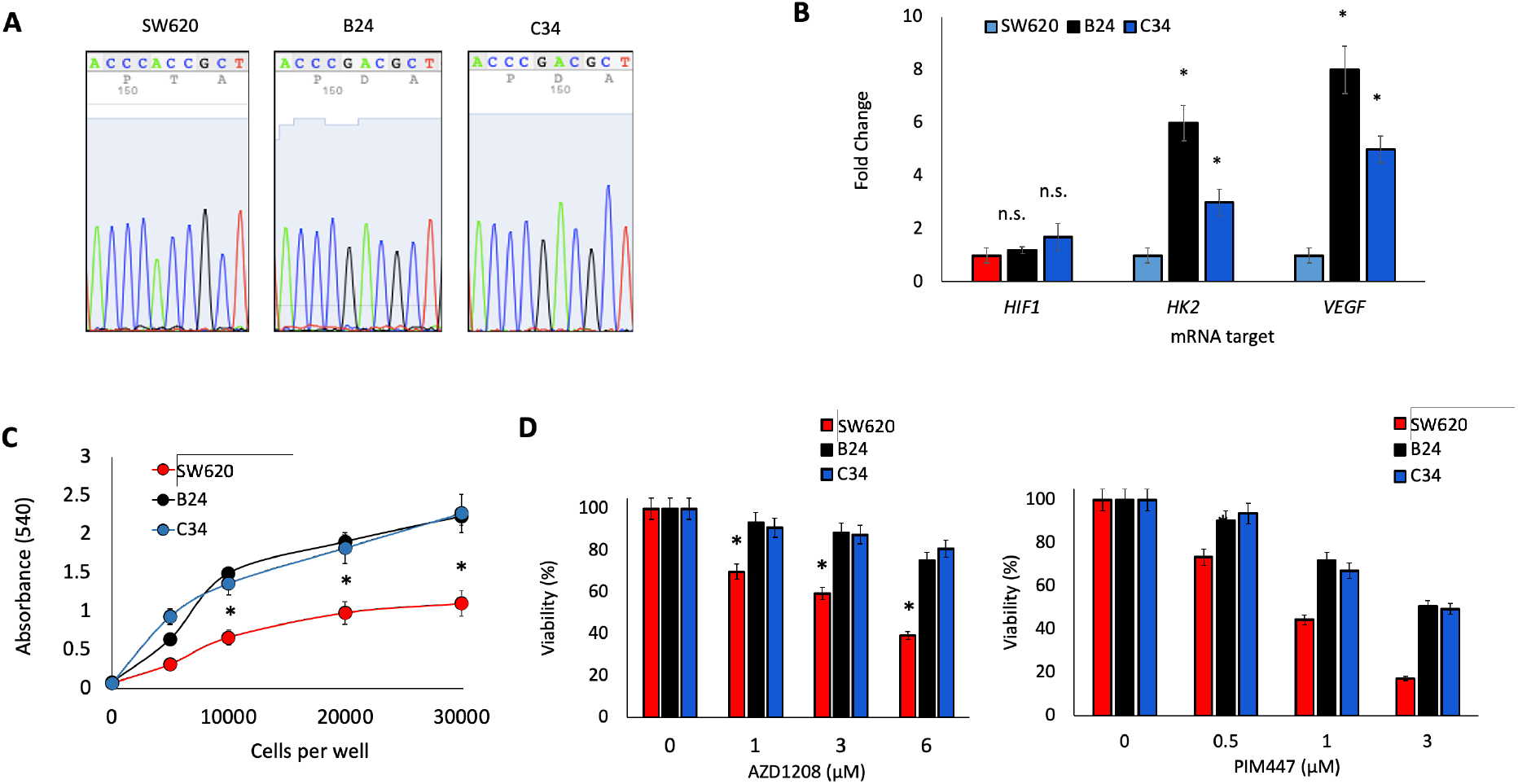
Validation and characterization of SW620 HIF-1α T455D CRISPR clones. A) Sanger sequencing spectra from SW620 parental and CRISPR clones. B) RNA was harvested from each cell line for qRT-PCR analysis. C) The indicated number of SW620, B24, and C34 cells were plated and allowed to grow for 48 h before analysis by MTT. D) Twenty thousand SW620, B24 and C34 cells were treated with various concentrations AZD1208 or PIM447 for 24 h before analysis by MTT. *p < 0.05, error bars = SEM.

**Supplemental Figure 5.**
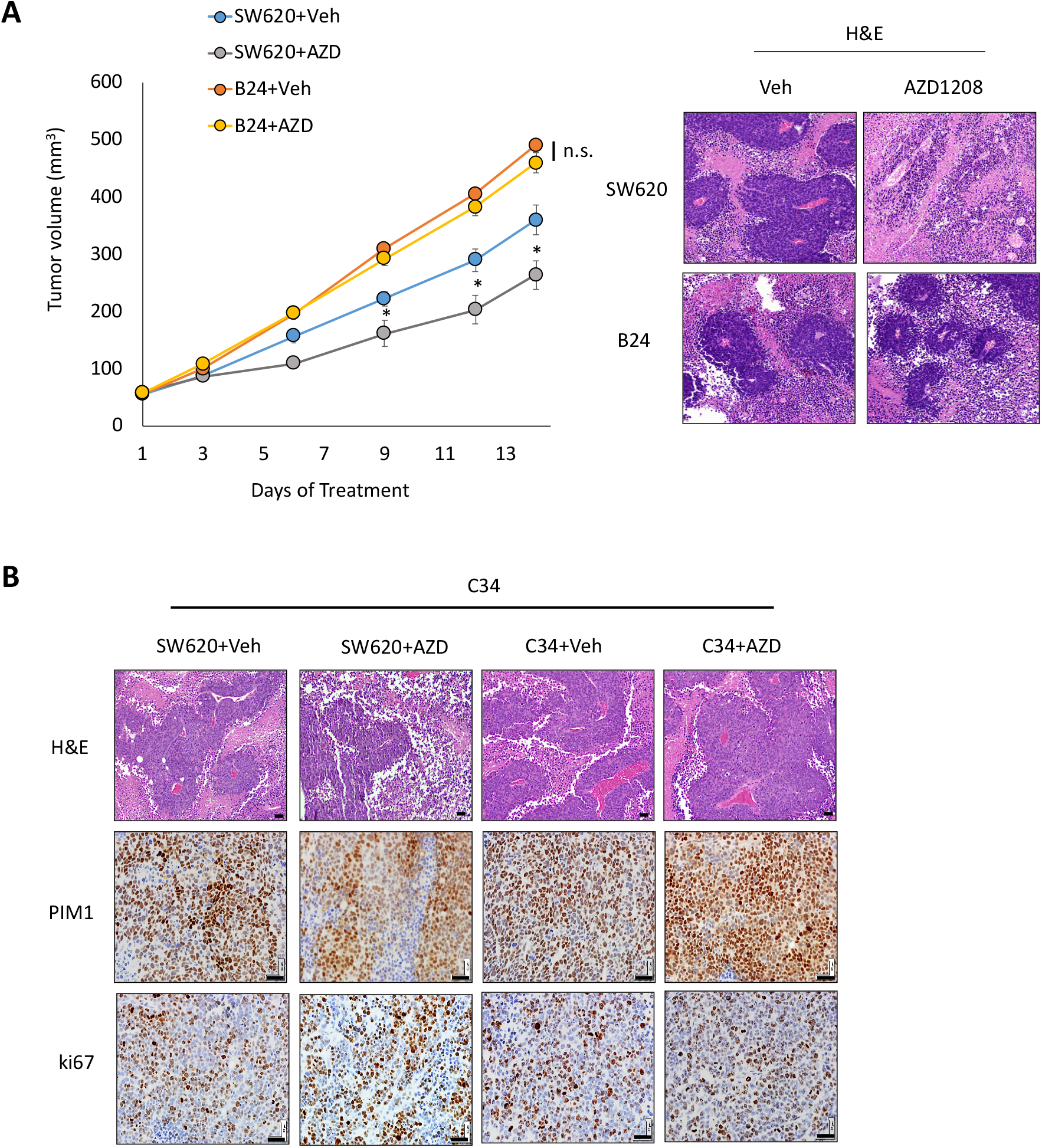
HIF-1α T455D mutant tumors are refractory to PIM inhibitor. A) Mice were injected 5×10^6^ SW620 or B24 cells, treated with vehicle or AZD1208 (30 mg/kg), and tumor volume was measured over time. Tumors from each cohort were stained for H&E to assess vascularization. B) Immunostaining of SW620 and C34 tumors after the indicated treatment *p < 0.05, error bars =SEM.

